# Transient developmental increase of prefrontal activity alters network maturation and causes cognitive dysfunction in adult mice

**DOI:** 10.1101/558957

**Authors:** Sebastian H. Bitzenhofer, Jastyn A. Pöpplau, Mattia Chini, Annette Marquardt, Ileana L. Hanganu-Opatz

**Author notes:** Co-first authors (Sebastian H. Bitzenhofer, Jastyn A. Pöpplau). Center for Neural Circuits and Behavior, Department of Neurosciences, University of California, San Diego, La Jolla, CA, USA. Corresponding authors: Ileana L. Hanganu-Opatz, Falkenried 94, 20251 Hamburg, Germany, Sebastian Bitzenhofer, 9500 Gilman Dr., La Jolla, CA 92093, USA.

## Abstract

Disturbed neuronal activity in neuropsychiatric pathologies emerges during development and might cause multifold neuronal dysfunction by interfering with apoptosis, dendritic growth and synapse formation. However, how altered electrical activity early in life impacts neuronal function and behavior of adults is unknown. Here, we address this question by transiently increasing the coordinated activity of layer 2/3 pyramidal neurons in the medial prefrontal cortex of neonatal mice and monitoring long-term functional and behavioral consequences. We show that increased activity during early development causes premature maturation of pyramidal neurons and alters interneuron density. Consequently, reduced inhibitory feedback by fast-spiking interneurons and excitation/inhibition imbalance in prefrontal circuits of young adults result in weaker evoked synchronization in gamma frequency. These structural and functional changes ultimately lead to poorer mnemonic and social abilities. Thus, prefrontal activity during early development actively controls the cognitive performance of adults and might be critical for cognitive symptoms of neuropsychiatric diseases.

## Main

The prefrontal cortex acts as a hub of cognitive processing indispensable for the daily life^1,2^. Disruption of prefrontal-dependent short-term memory and executive performance is the major burden of neuropsychiatric diseases, such as schizophrenia and autism spectrum disorders^3–5^. These diseases have been associated with a large variety of genes and environmental risk factors that increase susceptibility^6,7^. The absence of a clear understanding of their pathophysiology has resulted in primarily symptom-based treatments with low response rates^8^. Many of the genes and risk factors associated with neuropsychiatric diseases regulate brain development, leading to the hypothesis that abnormal maturation causes impaired network function and ultimately poor cognitive abilities later in life^8–11^. Indeed, rhythmic network activity of cortical, and particularly prefrontal circuits is already compromised in prodromal patients^12,13^ and during early postnatal development in mouse models of schizophrenia and autism^14–17^.

Neuronal activity regulates the development of cortical networks in many ways, from controlling neuronal differentiation, migration and apoptosis up to shaping the establishment of sensory maps, local and large-scale networks^18–21^. Early in life, activity in the prefrontal cortex is coordinated in oscillatory patterns, yet, in line with the delayed structural maturation and emergence of cognitive abilities, they appear later than in other cortical areas^22^. Inputs from cortical and subcortical areas boost the activation of local prefrontal circuits^22–25^. Moreover, intracortical interactions lead to the emergence of oscillatory activity at fast frequencies^26,27^. However, whether early activity is necessary for the maturation of prefrontal function and cognitive abilities is still unknown. Conversely, to which extent altered activity during development actively contributes to adult miswiring relevant for disease conditions, instead of simply reflecting pathological maturation, remains to be elucidated.

To address these questions, we manipulated cortical activity during early development and monitored the long-term consequences for network activity and behavioral abilities. The manipulation was achieved by transient light stimulation of a subset of pyramidal neurons (PYRs) in layer (L) 2/3 of the mouse medial prefrontal cortex (mPFC) from postnatal day (P) 7 to 11, the developmental time window corresponding to the second/third trimester of gestation in humans^28^. This light stimulation induces rhythmic activity in beta/low gamma frequency in the developing prefrontal cortex^26^. At the age of stimulation, the migration of cortical neurons has finished and the activity-dependent formation of synaptic connections is in full progress^18,29,30^. We focused on this critical developmental period for cortical network formation at which mouse models of psychiatric diseases start to show altered prefrontal activity caused by L2/3 PYRs dysfunction^14^. We demonstrate that the transient increase of prefrontal activity during early development is sufficient to disrupt prefrontal function and cognitive performance at young adult age.

## Results

### Stimulation of L2/3 pyramidal neurons induces coordinated activity in the neonatal mPFC

To uncover the role of early activity for adult prefrontal function, we firstly established a protocol to optically manipulate the activity of L2/3 PYRs from P7-11, the developmental time window critical for the formation of synaptic contacts in mPFC (Fig. 1a). For this, a subset of precursor cells of L2/3 PYRs in the prelimbic subdivision of the mPFC was transfected with channelrhodopsin 2 E123T/T159C (ChR2(ET/TC)) by in utero electroporation (IUE) at embryonic day (E) 15.5. As previously reported, the IUE protocol yields unilateral expression of ChR2(ET/TC) in 20-30% of PYRs confined to L2/3 in the mPFC (Fig. 1b)^31^.

**Fig. 1.**
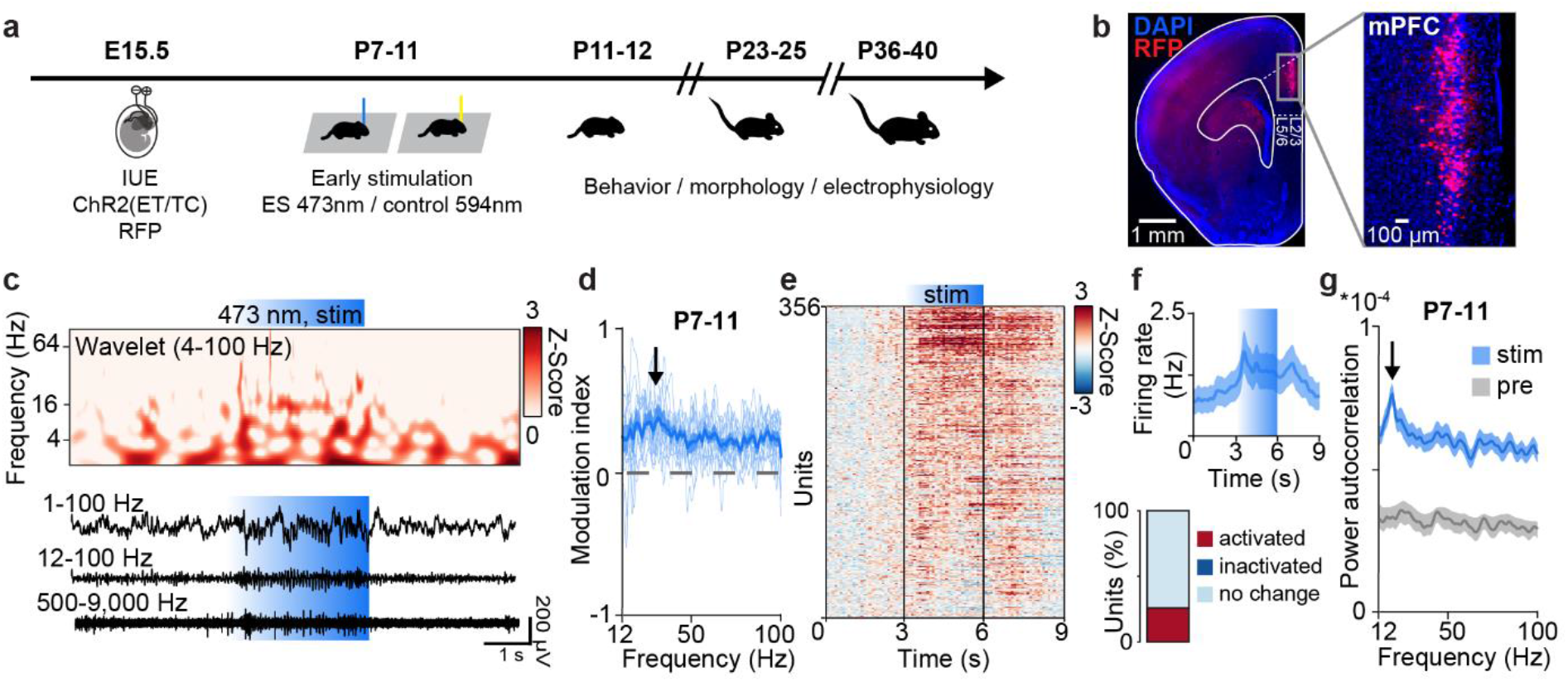
Light stimulation of L2/3 PYRs in the neonatal mPFC. (**a**) Schematic of the protocol for early light stimulation and long-lasting monitoring of structural, functional, and behavioral effects during development. (**b**) Representative image showing ChR2(ET/TC)-2A-RFP-expression in L2/3 PYRs after IUE at E15.5 in a DAPI-stained coronal slice including the mPFC from a P11 mouse. (**c**) Representative extracellular recording displayed together with corresponding wavelet spectrum at identical time scale during ramp light stimulation (473 nm, 3 s) of L2/3 PYRs in the mPFC of a P11 mouse. (**d**) Modulation index of local field potential (LFP) power in response to ramp light stimulation averaged for P7-11 mice (n=13). (**e**) Firing rates of single units (n=356 units from 13 mice) in response to ramp light stimulation z-scored to pre-stimulation period. (**f**) Single unit firing rate during ramp light stimulation averaged for P7-11 mice (top, n=356 units from 13 mice) and percent of significantly modulated units (bottom). (**g**) Power of single unit autocorrelations before (pre) and during (stim) ramp light stimulation averaged for P7-11 mice (n=356 units from 13 mice).

Ramp stimulations of linearly increasing light power (473 nm, 3 s) were used to activate transfected L2/3 PYRs from P7 to P11. In line with previous data^26^, prefrontal network activity tended to organize itself rhythmically at 15-20 Hz upon ramp stimulation (Fig. 1c,d). This rhythmic activity resembled the discontinuous activity spontaneously occurring in the neonatal mPFC^22,32^. Ramp light stimulation increased neuronal firing in a subset of neurons (20.2% of units significantly activated, 0.6% of units significantly inactivated) (Fig. 1e, f). Induced firing was not random, but peaked at 15-20 Hz for individual units, similar to induced network activity (Fig. 1g). Due to the thin skull at this age, similar activity was induced with transcranial light stimulation (Extended Data Fig. 1a). Control light stimulations (594 nm, ramp, 3 s) that do not activate ChR2(ET/TC) did not change the firing and network activity in the mPFC (Extended Data Fig. 1b-f).

**Extended Data Fig. 1.**
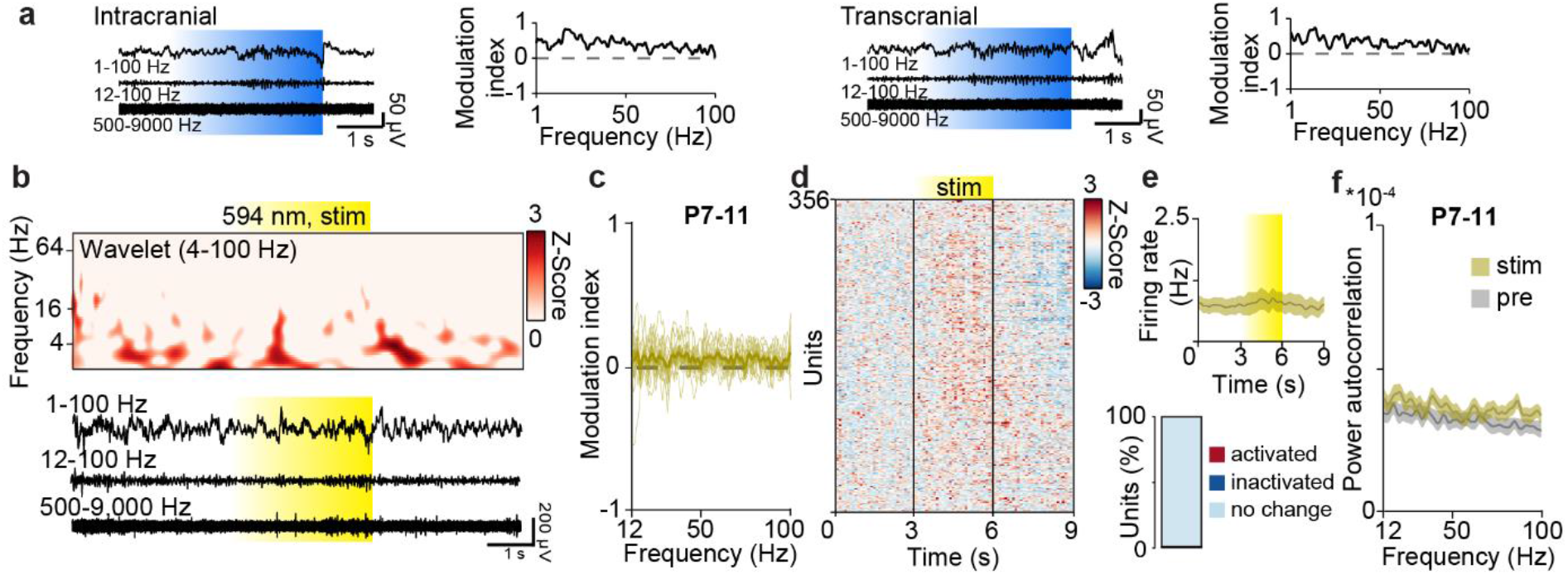
Control light stimulation of L2/3 PYRs in the neonatal mPFC. (**a**) Representative extracellular recordings during intracranial (left) and transcranial (right) ramp light stimulations (473 nm, 3 s) of L2/3 PYRs, as well as corresponding MI of power spectra for a P11 mouse. (**b**) Representative extracellular recording displayed together with corresponding wavelet spectrum at identical time scale during control ramp light stimulation (594 nm, 3 s) of L2/3 PYRs in the mPFC of a P11 mouse. (**c**) Modulation index of LFP power in response to control ramp light stimulation averaged for P7-11 mice (n=13). (**d**) Firing rates of single units (n=356 units from 13 mice) in response to control ramp light stimulation z-scored to pre-stimulation period. (**e**) Single unit firing rate during control ramp light stimulation averaged for P7-11 mice (top, n=356 units from 13 mice) and percent of significantly modulated units (bottom). (**f**) Power of single unit autocorrelations before (pre) and during (stim) control ramp light stimulation averaged for P7-11 mice (n=356 units from 13 mice).

### Transient increase of prefrontal activity during neonatal development disrupts cognitive performance of young adults

To transiently increase neuronal firing and network activation in the developing mPFC, we performed the transcranial stimulation that induced fast oscillatory discharges daily from P7 to P11. This developmental period has been identified as being critical for altered prefrontal activity in a mouse model of neuropsychiatric diseases^14^. On each of the five days of manipulation, mice received 30 transcranial ramp light stimulations (3 s long) at either 594 nm (control) or 473 nm (early stimulation, ES) to activate the ChR2(ET/TC)-transfected L2/3 PYRs in the mPFC.

Subsequently, we tested the behavioral abilities of control and ES mice, focusing on tasks that require prefrontal function. Data from mice of both sexes were pooled, since their performance was comparable in all tasks (Extended Data Tab. 2). Transient early stimulation did not affect the overall somatic and reflex development (Extended Data Fig. 2). First, we monitored recognition memory as a form of short-term memory that emerges at pre-juvenile age (P16-22), as soon as sensory and motor abilities are fully mature^33^. In contrast to control mice, ES mice were not able to distinguish a novel from a familiar object (novel object recognition, NOR) as well as an object they more recently interacted with (recency recognition, RR) (Fig. 2a,b). However, group differences were not significant for NOR and RR. The novel position of an object (object location recognition, OLR) was distinguished by both control and ES mice (Extended Data Fig. 3b). In contrast to NOR and RR, OLR depends more on hippocampus than mPFC^34^. Social interactions were significantly impaired in pre-juvenile ES mice. Their preference for interaction with the dam-containing container over an empty container was significantly reduced compared to control mice (Fig. 2c).

**Fig. 2.**
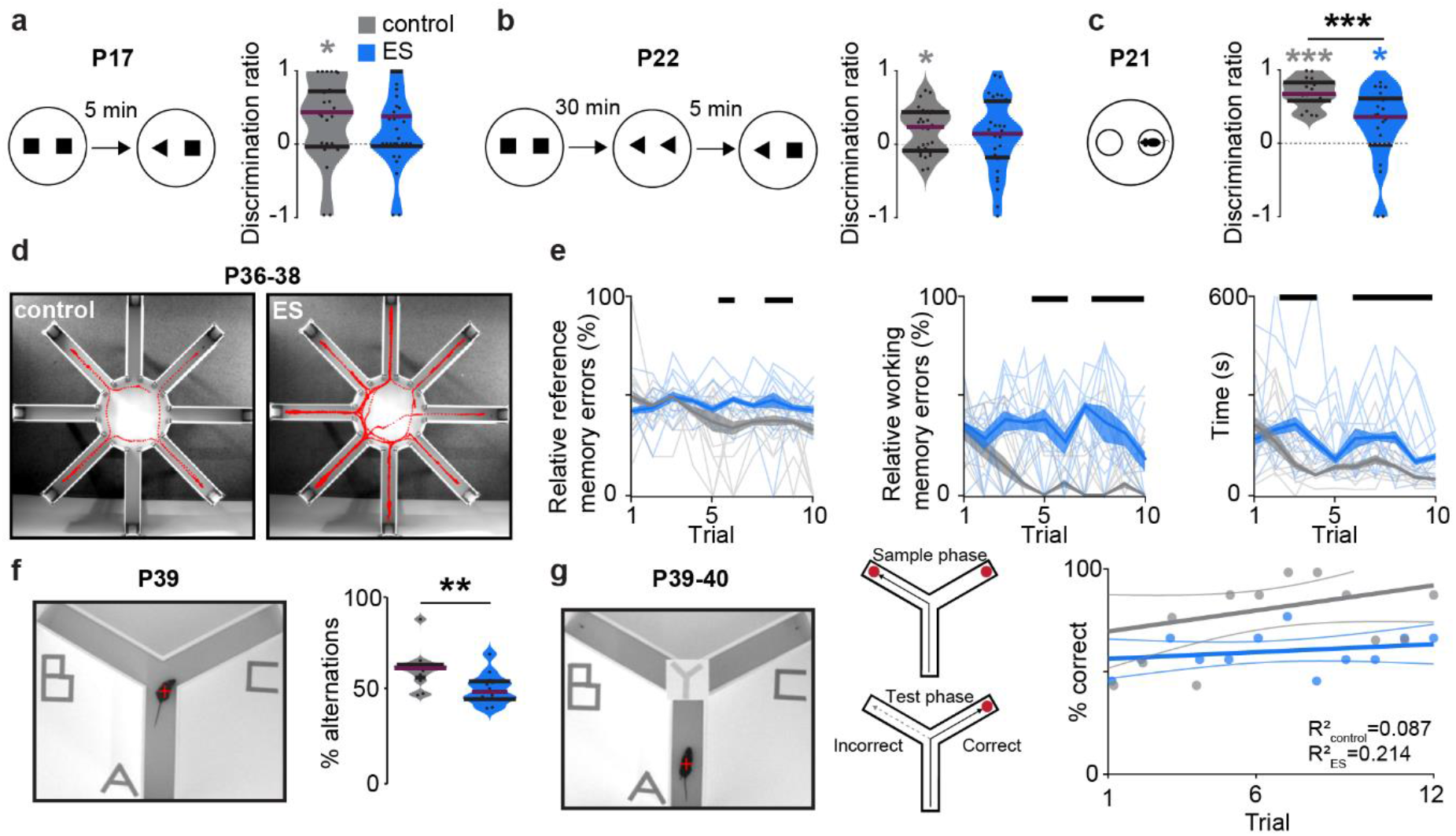
Transient early stimulation impairs cognitive abilities of juvenile and young adult mice. (**a**) Schematic of NOR task and violin plot displaying the discrimination ratio of interaction time with a novel vs. familiar object for control (n=28) and ES (n=30) mice at P17. (Wilcoxon rank, control p=0.018, ES p=0.157, control-ES p=0.177). (**b**) Schematic of RR task and violin plot displaying the discrimination ratio of interaction time with a less vs. more recent object for control (n=28) and ES (n=30) mice at P22. (Wilcoxon rank, control p=0.010, ES p=0.171, control-ES p=0.498). (**c**) Schematic of maternal interaction task and violin plot displaying the discrimination ratio of interaction time with mother vs. empty bin for control (n=19) and ES (n=21) mice at P21. (Wilcoxon rank, control p<0.001, ES p=0.045, control-ES p<0.001). (**d**) Representative tracking of a control (left) and ES mouse (right) in an 8-arm radial maze memory task with 4 baited arms at P36-38. (**e**) Plots displaying the relative reference (left) and working-memory errors (middle), as well as the time to complete the task (right) in 8-arm radial maze memory task over 10 trials on 3 consecutive days for control (n=12) and ES (n=12) mice. (Kruskall-Wallis, relative reference memory errors p<0.001, relative working memory errors p<0.001, time p<0.001). (**f**) Photograph illustrating a spontaneous alternation task in a Y-maze (left) and violin plot displaying the percent of spontaneous alternations (right) for control (n=12) and ES (n=12) mice at P39. (Wilcoxon rank, p=0.006). (**g**) Photograph illustrating a delayed non-match-to-sample task in a Y-maze (left) and dot plot displaying the percent of correct choices over 12 consecutive trials (6 trials/day) (right) for control (n=12) and ES (n=12) mice at P39-40. Black lines and asterisks (* p<0.05, ** p<0.01, *** p<0.001) indicate significant differences (see Extended Data Tab. 1 for detailed statistics).

Second, we tested mPFC-dependent working memory of young adult (P36-40) control and ES mice. To this end, we used an 8-arm radial maze test with 4 baited arms, a Y-maze test for spontaneous alternations and a delayed non-match-to-sample task. ES mice showed working memory and reference memory deficits in the 8-arm radial maze test (Fig. 2d,e, Extended Data Fig.3d). Moreover, when compared to controls, ES mice showed poorer performance during spontaneous alternation (Fig. 2f, Extended Data Fig.3c). and in the delayed non-match-to-sample task (Fig. 2g). The deficits identified in ES mice are not due to impaired motor abilities or enhanced anxiety, since neither the behavior in an open field nor the interaction with objects and mazes was different between groups (Extended Data Fig. 3a,c,d). Thus, transient elevation of prefrontal activity at neonatal age caused long-lasting impairment of mPFC-dependent short-term and working memory as well as social behavior.

**Extended Data Fig. 2.**
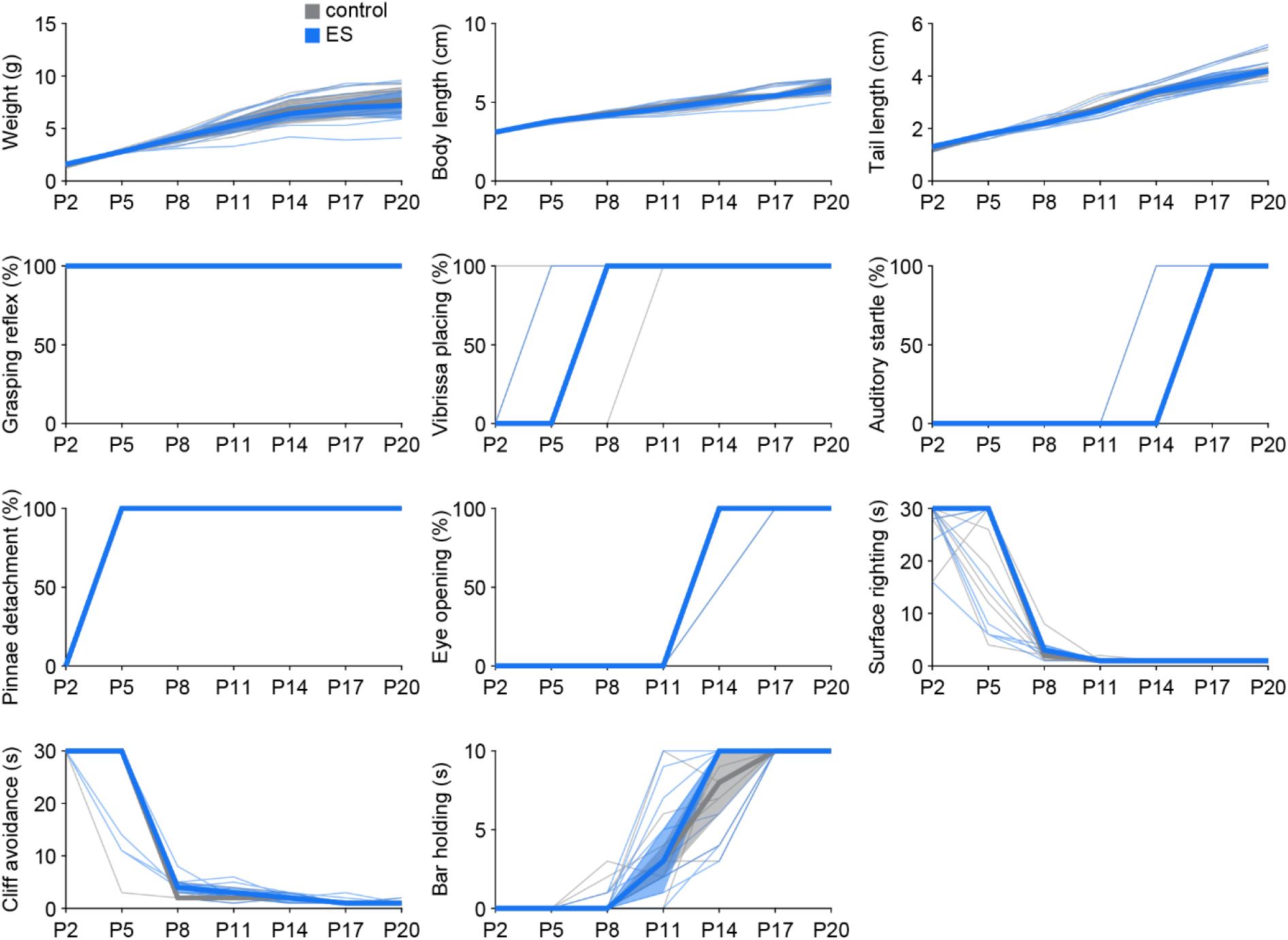
ES and control mice have similar somatic and reflex development. Line plots displaying the age-dependence of developmental milestones for control (n=11) and ES (n=11) mice. (See Extended Data Tab. 1 for detailed statistics).

**Extended Data Fig. 3.**
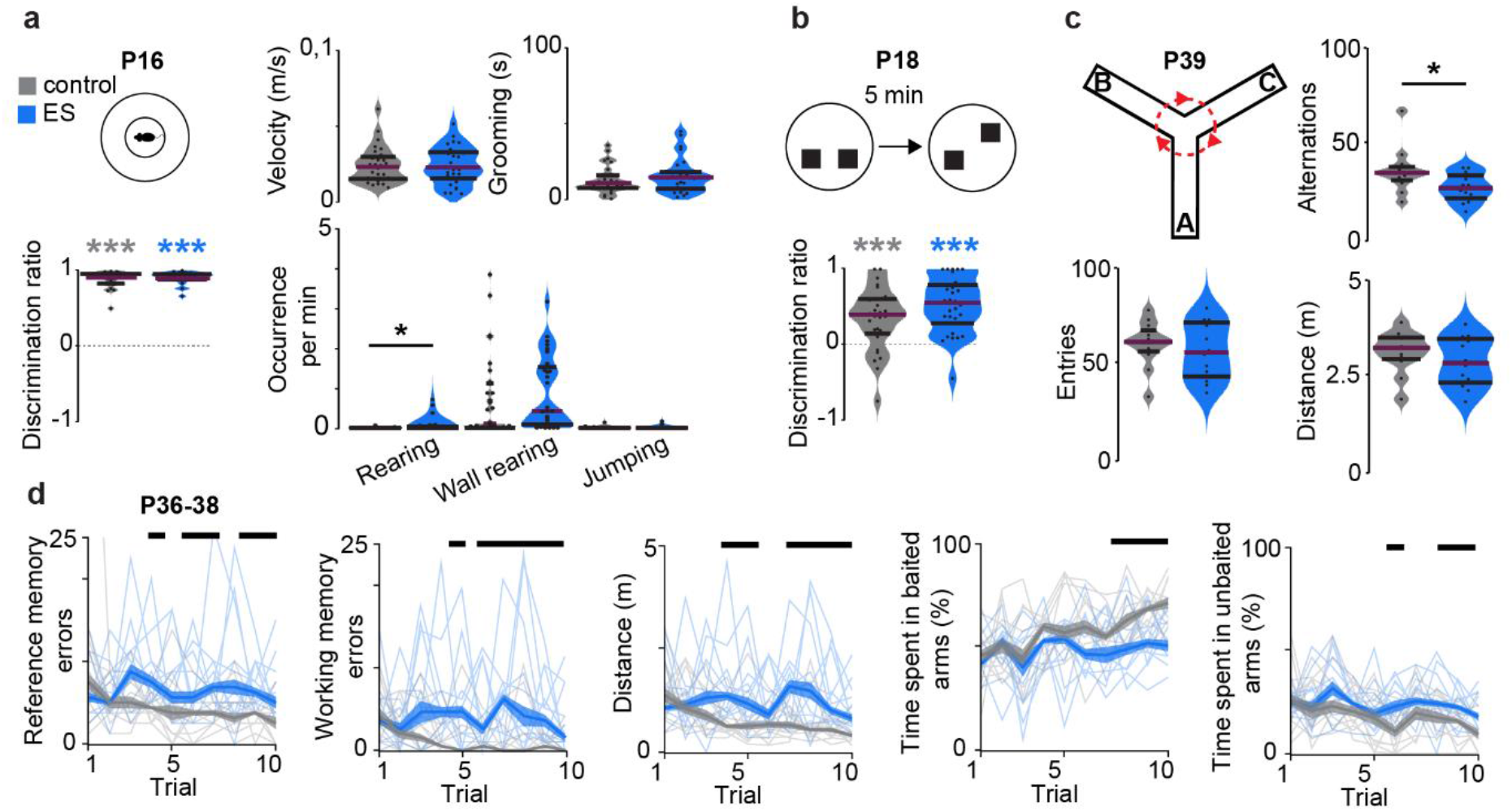
Transient early stimulation impairs mPFC-dependent cognitive abilities but not motor and anxiety behavior of juvenile and young adult mice. (**a**) Schematic of an open field task (top left) and violin plots displaying the discrimination ratio of time spend in border area vs. center area (bottom left), as well as the basic behavior (velocity, grooming, rearing, wall rearing, jumping) (right) for control (n=28) and ES (n=30) mice at P16. (Wilcoxon rank, discrimination ratio, control p<0.001, ES p<0.001, control-ES p=0.809). (**b**) Schematic of OLR task (top) and violin plot displaying the discrimination ratio of interaction time with an object in a novel vs. familiar location (bottom) for control (n=28) and ES (n=30) mice at P18. (Wilcoxon rank, control p<0.001, ES p<0.001, control-ES p=0.154). (**c**) Schematic showing spontaneous alternation in a Y-maze as well as violin plots displaying quantified parameters (alternations, entries, distance) for control (n=12) and ES (n=12) mice at P39. (Wilcoxon rank, alternations, p=0.046). (**d**) Line plots displaying reference- and working-memory errors as well as further task-related parameters for an 8-arm radial maze memory task over 10 trials on 3 consecutive days for control (n=12) and ES (n=12) mice at P36-38. (Kruskal-Wallis, reference memory errors p<0.001, working memory errors p<0.001). Black lines and asterisks (* p<0.05, ** p<0.01, *** p<0.001) indicate significant differences (see Extended Data Tab. 1 for detailed statistics).

### Transient increase of neonatal prefrontal activity induces premature dendritic growth in L2/3 pyramidal neurons

To test whether impaired cognitive abilities of juvenile and adult ES mice resulted from permanent structural disruption of the mPFC after transient increase of neonatal activity, we monitored the structural maturation of PYRs in control and ES mice. The density of CaMKII-positive neurons and of ChR2(ET/TC)-transfected neurons did not differ between control and ES mice at all investigated developmental time points (P11-12, P23-25 and P38-40) (Extended Data Fig. 4). Investigation of the dendritic morphology of L2/3 PYRs after transient stimulation at P7-11 revealed that immediately after this time window the dendritic arborization (i.e. dendrite length, number of intersections) of these neurons was increased in ES compared to control mice (Fig. 3). However, the exuberant arborization was transient and from P23-25 on, the dendritic arbors of L2/3 PYRs in the mPFC of ES mice were similar to controls. A comparison across age revealed that the dendritic length increased with age for control (linear mixed effect models (LMEM), P11-12 to P23-25 p=0.002**, P11-12 to P38-40 p=0.0002***), but not for ES mice (LMEM, P11-12 to P23-25 p=0.79, P11-12 to P38-40 p=0.07). Of note, dendritic length of L2/3 PYRs in ES mice at P11-12 was comparable to control mice at P23-25 (LMEM, p=0.33). These results suggest that increased activity in the neonatal mPFC causes premature dendritic maturation of L2/3 PYRs.

**Fig. 3.**
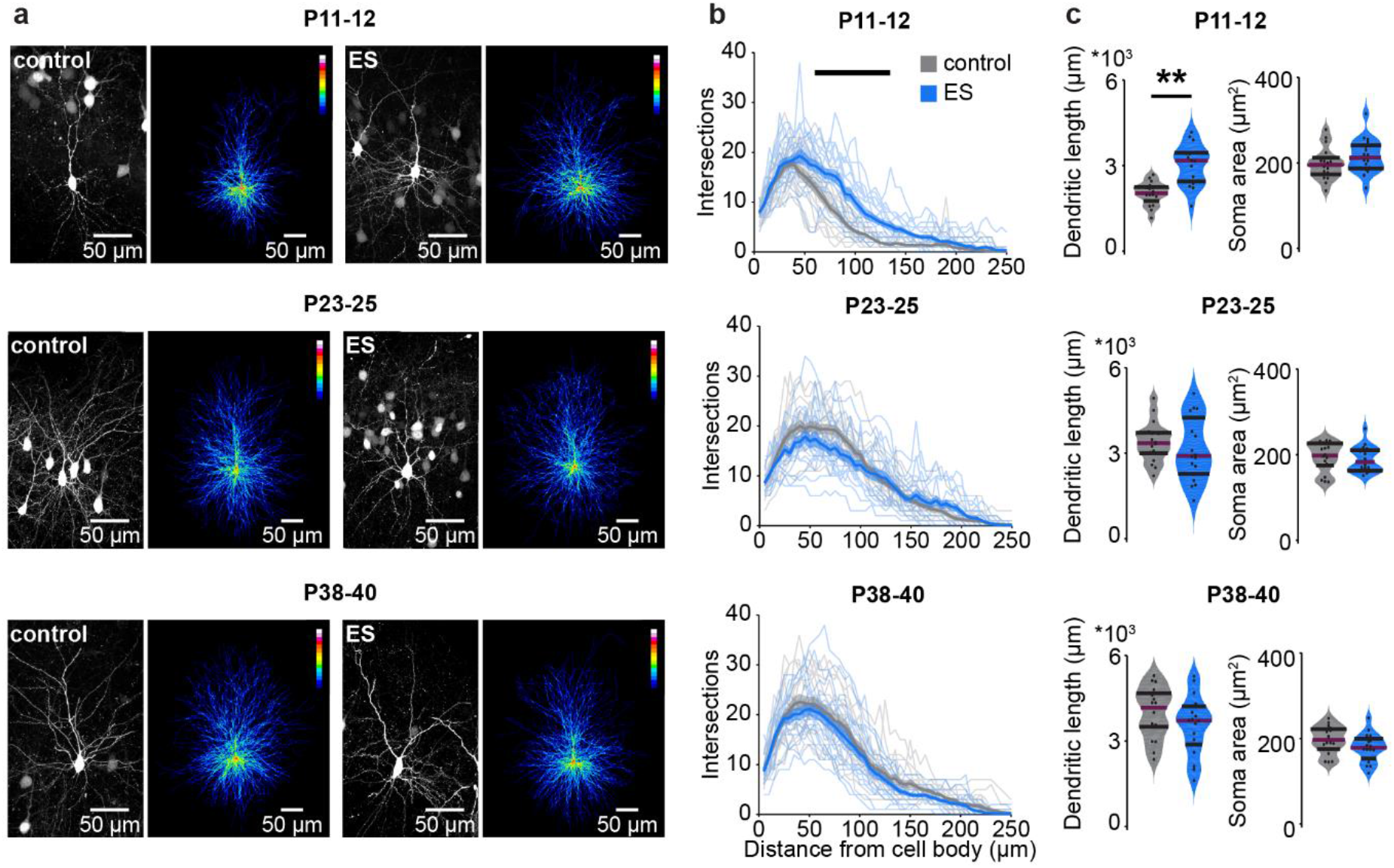
Transient early stimulation induces premature dendritic growth in prefrontal L2/3 PYRs. (**a**) Representative photographs and corresponding average heat maps of ChR2(ET/TC)-transfected L2/3 PYRs in the mPFC of P11-12, P23-25 and P38-40 control (left) and ES mice (right). (**b**) Line plots of dendritic intersections of L2/3 PYRs with concentric circles (0-250 μm radius) centered around the soma averaged for control (18 cells of 3 mice/age group) and ES mice (18 cells of 3 mice/age group) at P11-12, P23-25 and P38-40. (LMEM, P11-12 p<0.001, P23-25 p<0.001, P38-40 p<0.001). (**c**) Violin plots displaying the dendritic length and soma area of L2/3 PYRs for control (18 cells from 3 mice/age group) and ES (18 cells from 3 mice/age group) mice for different age groups. (LMEM, dendritic length, P11-12 p=0.007, P23-25 p=0.631, P38-40 p=0.161). Black lines and asterisks (* p<0.05, ** p<0.01, *** p<0.001) indicate significant differences (see Extended Data Tab. 1 for detailed statistics).

**Extended Data Fig. 4.**
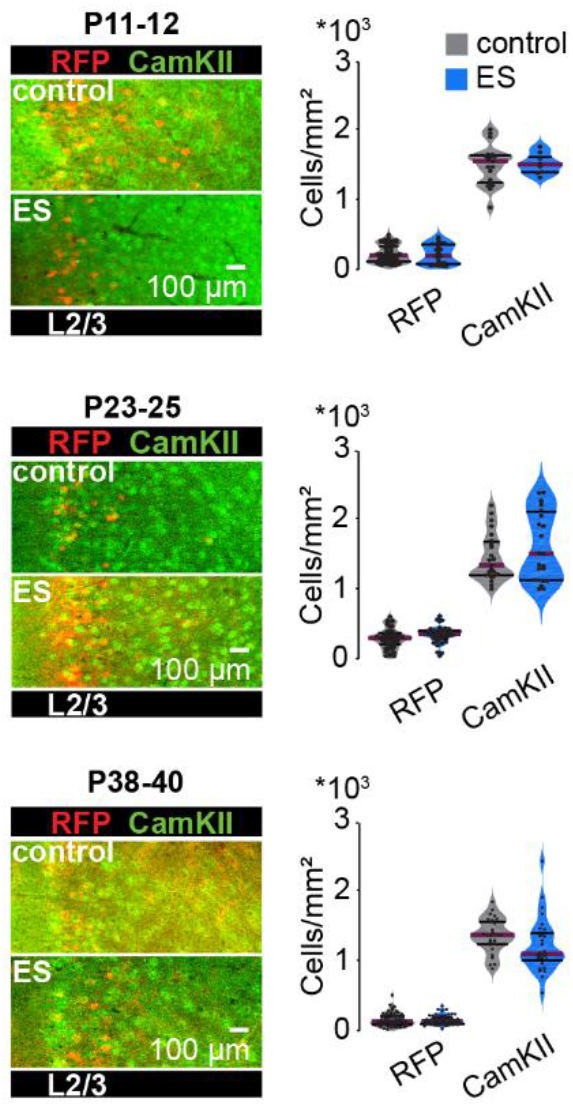
Transient early stimulation does not alter the density of L2/3 PYRs. Left, representative photographs displaying CaMKII immunostainings in the ChR2(ET/TC)-RFP-transfected mPFC of control and ES mice at P11-12 (control, RFP n=60 slices of 9 mice, CamKII n=19 slices of 6 mice; ES, RFP n=27 slices of 4 mice, CamKII n=9 slices of 2 mice), P23-25 (control, RFP n=47 slices of 5 mice, CamKII n=23 slices of 5 mice; ES, RFP n=43 slices of 5 mice, CamKII n=23 slices of 5 mice) and P38-40 (control, RFP n=65 slices of 5 mice, CamKII n=29 slices of 5 mice; ES, RFP n=62 slices of 5 mice, CamKII n=29 slices of 5 mice). Right, violin plots of RFP-expressing and CaMKII-positive neuronal density at different age groups. (LMEM, P11-12, RFP p=0.855, CamKII p=0.705, P23-25, RFP p=0.819, CamKII p=0.527, P38-40, RFP p=0.819, CamKII p=0.177). (See Extended Data Tab. 1 for detailed statistics).

### Transient increase of neonatal prefrontal activity reduces gamma power and network synchrony in the adult mPFC

Transient alteration of neonatal activity might perturb the function of prefrontal circuits, ultimately leading to abnormal behavior. To test this hypothesis, we monitored spontaneous neuronal and network activity of the mPFC across development. We performed extracellular recordings from head-fixed control and ES mice immediately after transient early stimulation (P11-12), at juvenile (P23-25) and young adult (P38-40) age (Fig. 4a,b). With increasing age, spontaneous oscillatory activity in the mPFC of control and ES mice increased in power and fast oscillations within 12-100 Hz became more prominent (Fig. 4c). At P11-12, the power of these fast oscillations was increased in the mPFC of ES mice compared to control mice, in accordance with the premature growth of L2/3 PYRs dendrites. At later stages of development, no differences were detected between control and ES mice. In contrast, the firing rates of single units were similar in control and ES mice during development, yet, at adulthood, ES mice showed decreased firing in the mPFC (Fig. 4d).

**Fig. 4.**
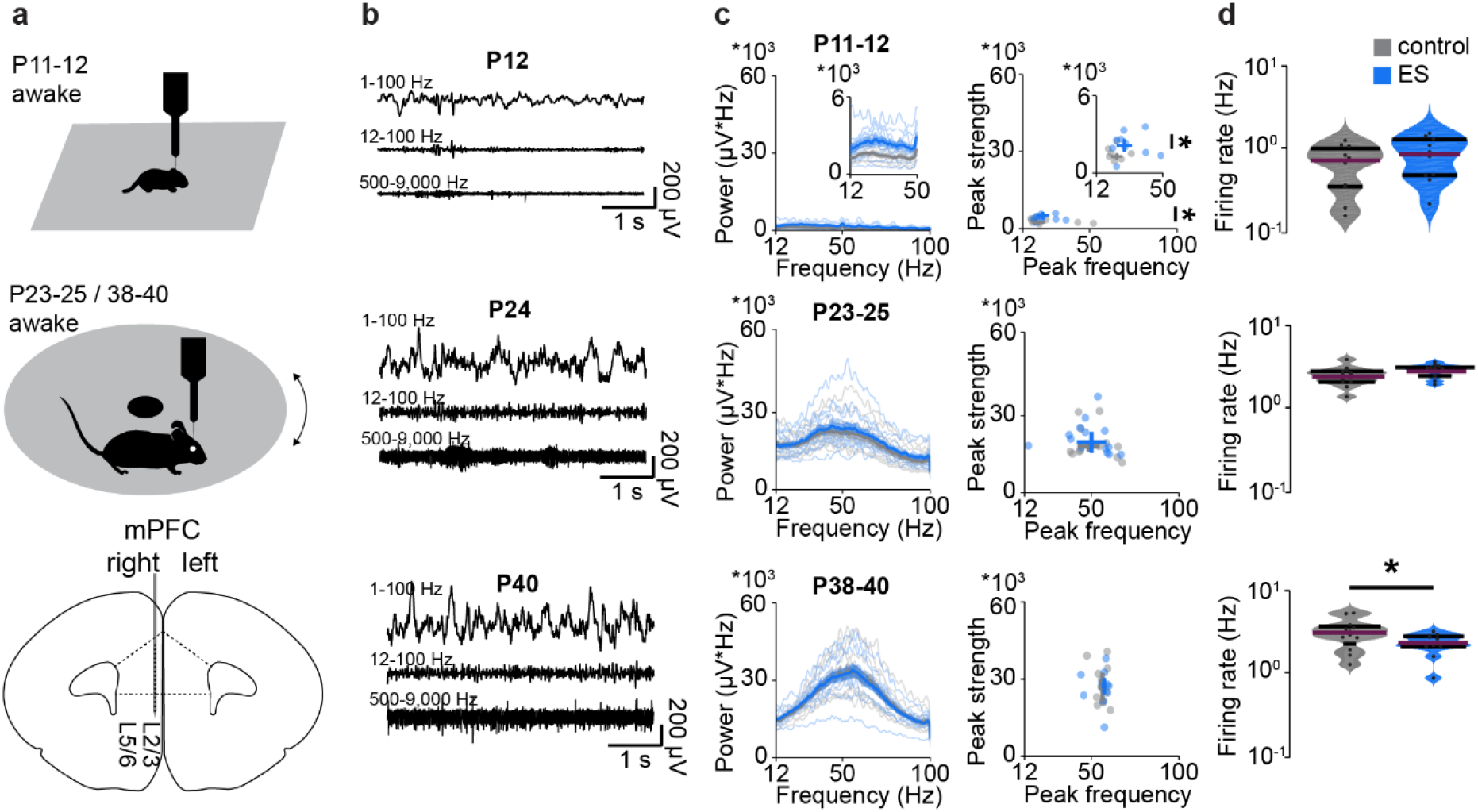
Transient early stimulation has minor effects on spontaneous network activity and firing in the developing mPFC. (**a**) Top, schematic illustrating the recording setups used for young mice with limited motor abilities (head-fixed, no movement) and for juvenile and young adult mice (head-fixed, freely moving on a spinning disk). Bottom, schematic of the recording configuration in the developing mPFC. (**b**) Representative extracellular recordings in the mPFC at P12, P24 and P40. (**c**) Left, average power spectra of spontaneous network activity in the mPFC of control and ES mice at P11-12 (control n=11 recordings, 11 mice, ES n=10 recordings, 10 mice), P23-25 (control n=13 recordings, 6 mice, ES n=14 recordings, 5 mice) and P38-40 (control n=12 recordings, 5 mice, ES n=12 recordings, 5 mice). Inset, power spectra for P11-12 shown at higher magnification. Right, scatter plots displaying peak strength and peak frequency of LFP power for control and ES mice. (Wilcoxon rank, P11-12, peak frequency p=0.245, peak strength p=0.015, LMEM, P23-25, peak frequency p=0.643, peak strength p=0.665, P38-40, peak frequency p=0.856, peak strength p=0.750). (**d**) Violin plots displaying the firing rates of single units in the mPFC averaged for control and ES mice at P11-12, P23-25, and P38-40. (Wilcoxon rank, P11-12 p=0.275, LMEM, P23-25 p=0.072, P38-40 p=0.041). Asterisks (* p<0.05, ** p<0.01, *** p<0.001) indicate significant differences (see Extended Data Tab. 1 for detailed statistics).

Even though spontaneous activity is largely unaffected by the transient increase of activity at neonatal age, the mPFC might abnormally respond to incoming stimuli, leading to disrupted processing and ultimately, behavior. To test this hypothesis, we used optogenetics to stimulate ChR2(ET/TC)-transfected L2/3 PYRs. Acute light stimulations (ramp, 473 nm, 3 s) triggered fast rhythmic activity with peak frequencies increasing from 15-20 Hz (beta frequency range) at P11-12 to 50-60 Hz (gamma frequency range) at P23-25 and P38-40 in the mPFC of control and ES mice (Fig. 5a). These results are in line with recent data, showing an acceleration of fast frequency oscillations during prefrontal development^35^. However, at P38-40 the magnitude of light-induced gamma activity was significantly smaller in ES mice compared to controls. This weaker prefrontal activation in fast oscillatory rhythms upon acute stimulation for ES mice, specific for young adults, was replicated in a separate cohort of anesthetized head-fixed mice (Extended Data Fig. 5a-c). Furthermore, young adult ES mice had weaker synchrony within and between hemispheres during evoked activity. Both, the coherence between L2/3 and L5/6 of the stimulated hemisphere and the coherence between L2/3 across hemispheres was reduced in ES mice at P38-40, but was normal at younger age (Extended Data Fig. 5d,e).

**Fig. 5.**
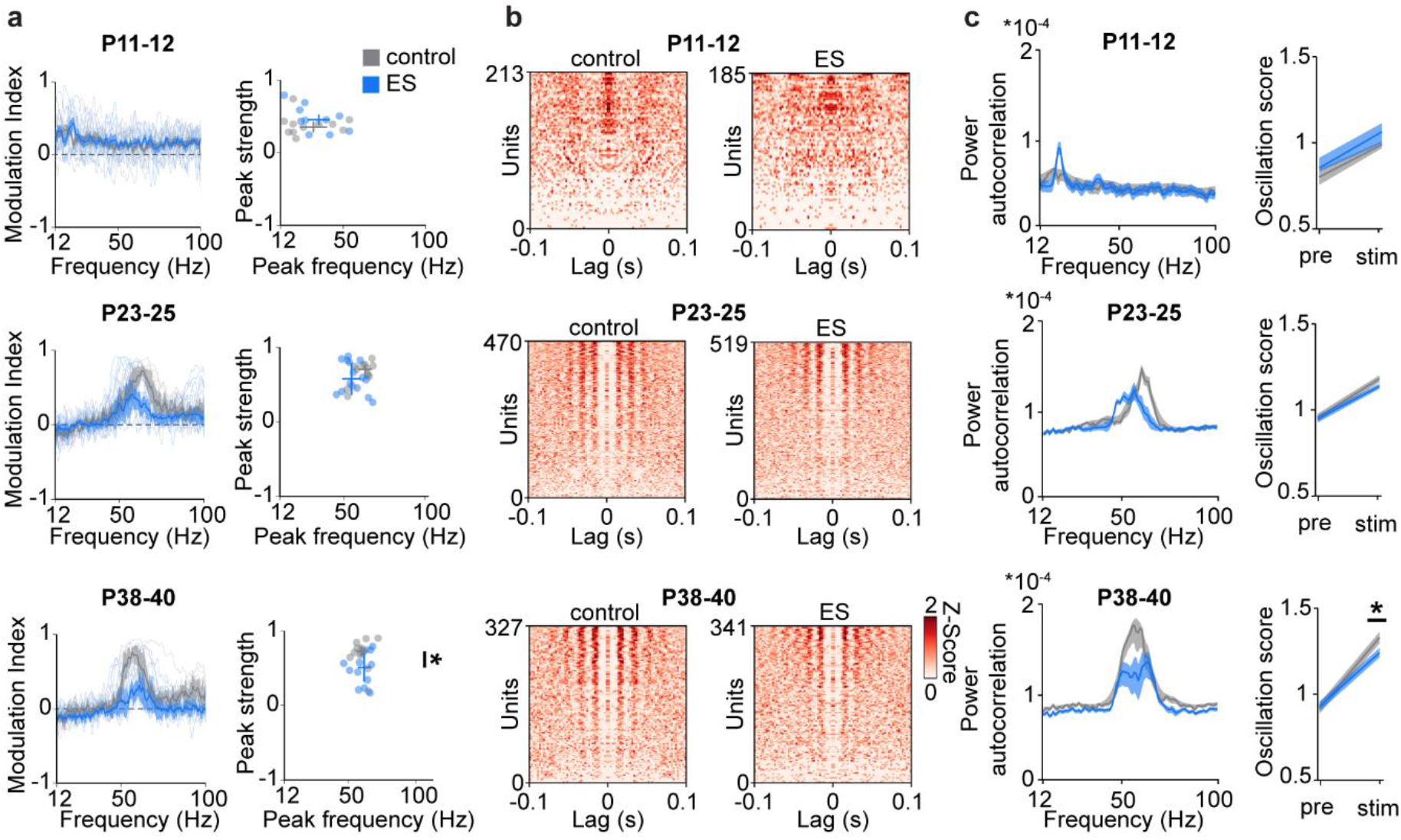
Transient early stimulation decreases evoked network and neuronal gamma rhythmicity in the adult mPFC. (**a**) Left, modulation index of LFP power in response to acute ramp light stimulation (473 nm, 3 s) for control and ES mice at P11-12 (control n=11 recordings, 11 mice, ES n=10 recordings, 10 mice), P23-25 (control n=13 recordings, 6 mice, ES n=14 recordings, 15 mice) and P38-40 (control n=12 recordings, 5 mice, ES n=12 recordings, 5 mice). Right, scatter plots displaying the peak strength and peak frequency of the power modulation index for control and ES mice. (Wilcoxon rank, P11-12, peak frequency p=0.307, peak strength p=0.307, LMEM, P23-25, peak frequency p=0.136, peak strength p=0.419, P38-40, peak frequency p=0.913, peak strength p=0.043). (**b**) Z-scored autocorrelograms of single units during acute ramp light stimulation arranged by magnitude for control and ES mice at P11-12 (control n=213 units, 11 mice, ES n=185 units, 10 mice), P23-25 (control n=470 units, 6 mice, ES n=519 units, 5 mice) and P38-40 (control n=327 units, 5 mice, ES n=341 units, 5 mice). (**c**) Left, average power of single unit autocorrelograms during acute ramp light stimulation for control and ES mice at different age. Right, oscillation score of single units before (pre) and during (stim) acute ramp light stimulation. (LMEM, oscillation score, P11-12, pre p=0.406, stim p=0.156, P23-25, pre p=0.272, stim p=0.478, P38-40, pre p=0.428, stim p=0.030). Asterisks (* p<0.05, ** p<0.01, *** p<0.001) indicate significant differences (see Extended Data Tab. 1 for detailed statistics).

Additionally, we analyzed single unit firing to assess the response of prefrontal neurons to acute light stimulation in control and ES mice across development. Calculation of autocorrelations for prefrontal units showed that independent of age and group, neurons fire rhythmically in response to acute light stimulation (Fig. 5b). Similar to network oscillations, the strength and frequency of the rhythmicity of neuronal firing increased with age, yet the magnitude of increase was lower for ES mice, reaching significance at P38-40. In contrast, the rhythmicity of spontaneous firing of prefrontal units was similar for control and ES mice at all age groups (Extended Data Fig. 6).

**Extended Data Fig. 5.**
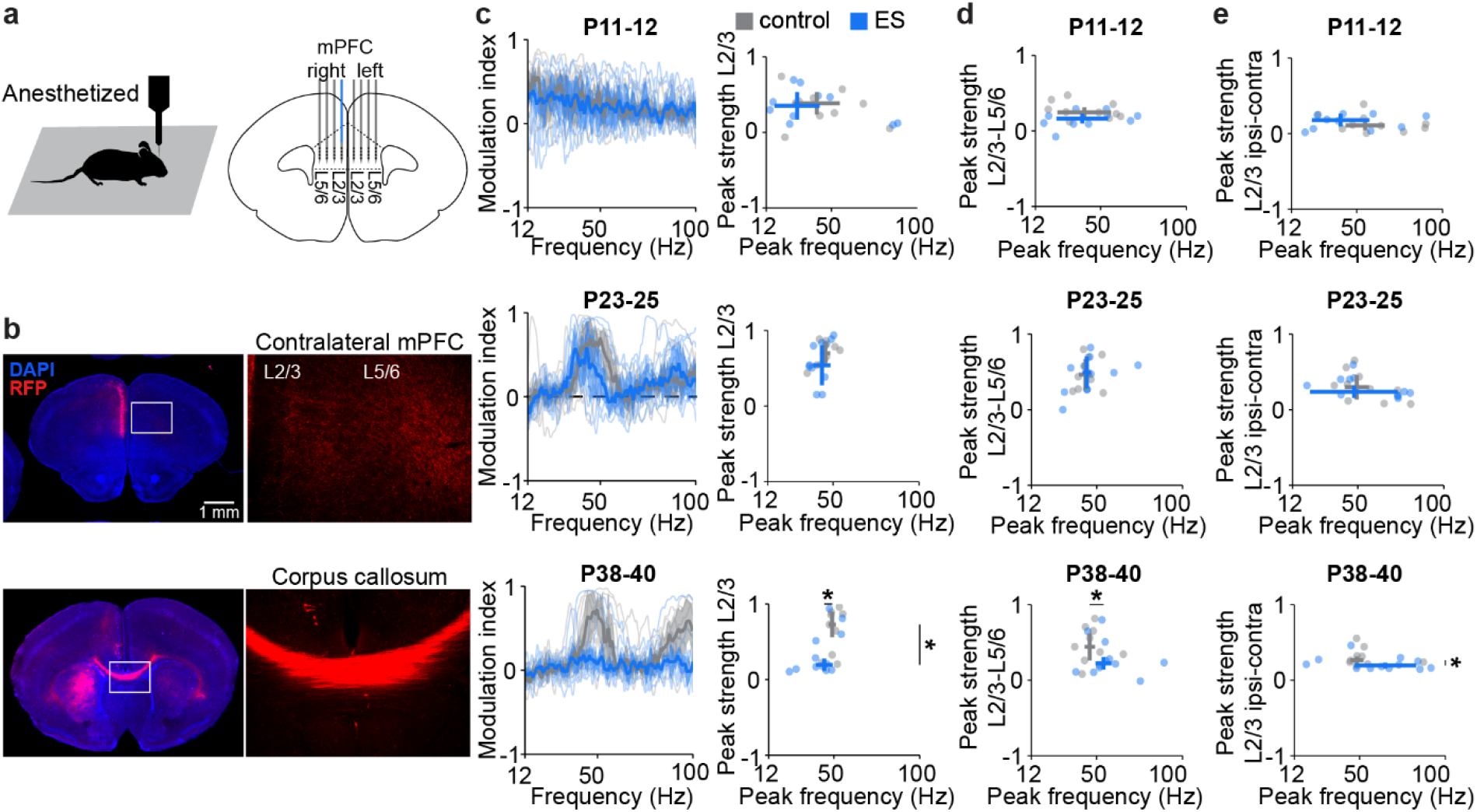
Transient early stimulation impairs evoked intra- and interhemispheric synchrony in the adult mPFC. (**a**) Schematic displaying bilateral multi-shank recordings in the mPFC of anesthetized mice. (**b**) Representative photographs showing axonal projections of ChR2(ET/TC)-RFP-transfected L2/3 PYRs in coronal slices from a P10 mouse. (**c**) Left, modulation index of LFP power in response to acute ramp light stimulation (473 nm, 3 s) for control and ES mice at P11-12 (control n=10 recordings, 10 mice, ES n=10 recordings, 10 mice), P23-25 (control n=10 recordings, 10 mice, ES n=11 recordings, 11 mice) and P38-40 (control n=9 recordings, 9 mice, ES n=12 recordings, 12 mice). Right, scatter plots displaying the peak strength and peak frequency of the power modulation index for control and ES mice. (Wilcoxon rank, P11-12, peak frequency p=0.520, peak strength p=0.909, P23-25, peak frequency p=0.290, peak strength p=0.459, P38-40, peak frequency p=0.039, peak strength p=0.025). (**d**) Scatter plots displaying the peak strength and peak frequency of prefrontal L2/3-L5/6 coherence at different age. (Wilcoxon rank, P11-12, peak frequency p=1.000, peak strength p=0.053, P23-25, peak frequency p=0.943, peak strength p=0.915, P38-40, peak frequency p=0.042, peak strength p=0.069). (**e**) Scatter plots displaying the peak strength and peak frequency of interhemispheric prefrontal L2/3-L2/3 coherence at different age. (Wilcoxon rank, P11-12, peak frequency p=0.212, peak strength p=0.623, P23-25, peak frequency p=0.832, peak strength p=0.915, P38-40, peak frequency p=0.270, peak strength p=0.036). Asterisks (* p<0.05, ** p<0.01, *** p<0.001) indicate significant differences (see Extended Data Tab. 1 for detailed statistics).

**Extended Data Fig. 6.**
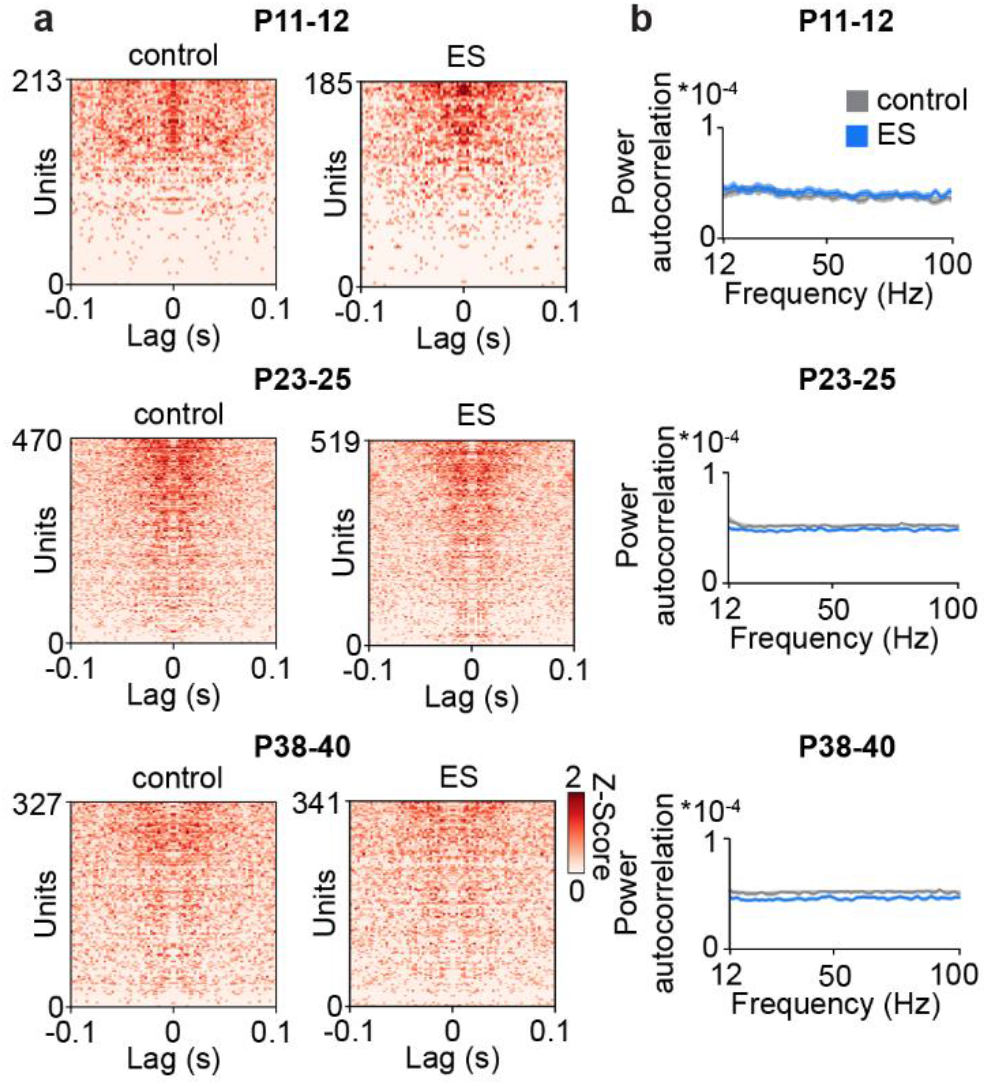
Transient early stimulation does not affect the rhythmicity of single units in the mPFC during spontaneous activity. (**a**) Z-scored autocorrelations for single units before acute ramp light stimulation arranged by magnitude for control and ES mice at P11-12 (control n=213 units, 11 mice, ES n=185 units, 10 mice), P23-25 (control n=470 units, 6 mice, ES n=519 units, 5 mice) and P38-40 (control n=327 units, 5 mice, ES n=341 units, 5 mice). (**b**) Average power of single unit autocorrelations before acute ramp light stimulation for control and ES mice at different age. (See Extended Data Tab. 1 for detailed statistics).

To assess the impact of early stimulation on the synchrony within the prefrontal network during development, we calculated pairwise correlations of single units. During spontaneous activity, pairwise correlations between prefrontal units were similar for control and ES mice at P11-12 and P38-40, whereas correlation at the 3^rd^ quartile was slightly reduced in ES mice at P23-25 (Extended data Fig. 7). In contrast, during ramp light stimulations, the pairwise correlations were significantly reduced at the 3^rd^ quartile in young adult ES mice when compared to controls, but comparable between groups at P11-12 and P23-25. These data show that the synchrony of the highest correlated units in the mPFC is reduced in young adult ES mice.

Taken together, these results show that transiently increased activity at neonatal age diminishes prefrontal gamma band synchronization in response to stimulation at adult age.

**Fig. 6.**
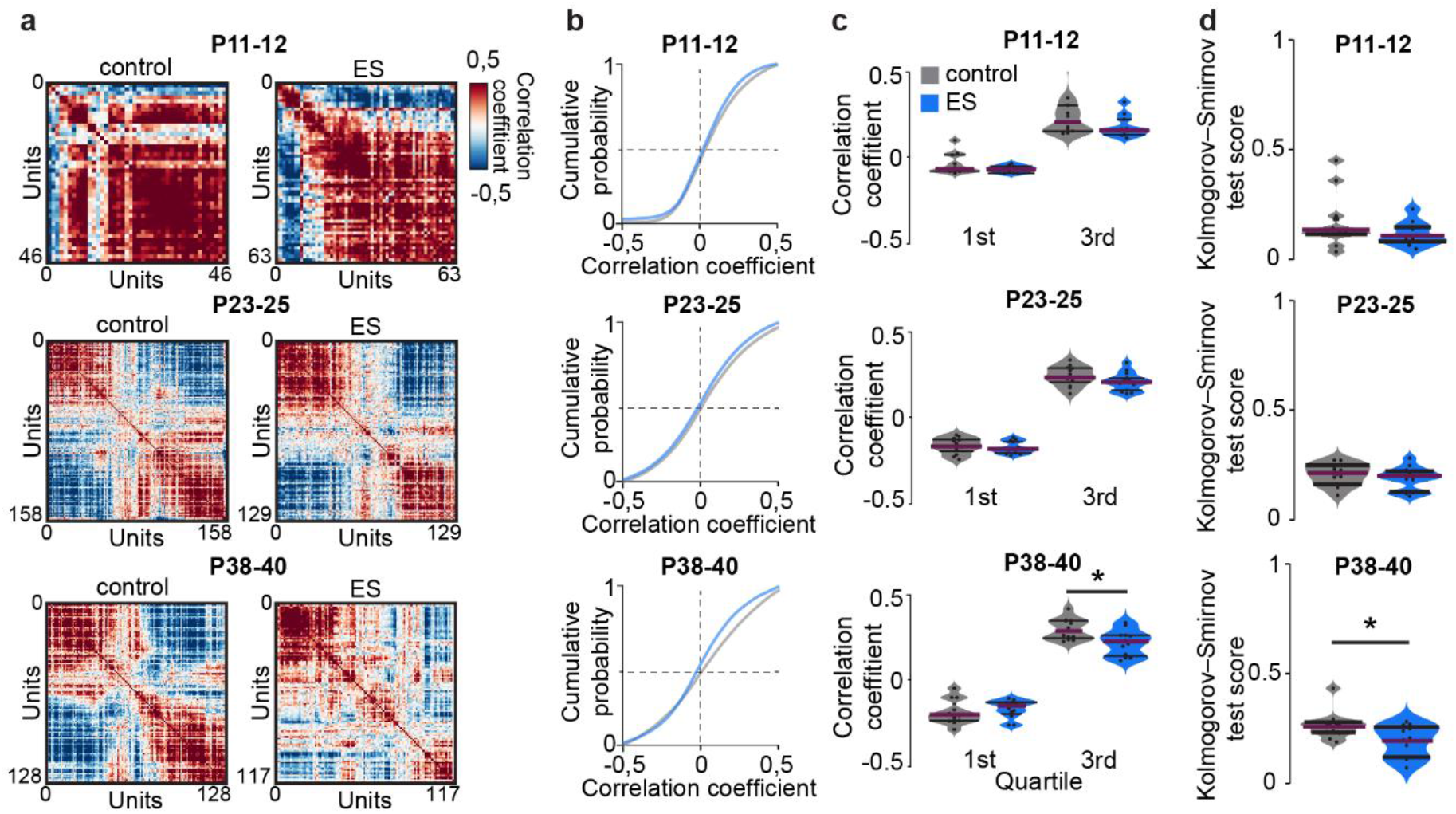
Transient early stimulation reduces synchrony of highly correlated units in response to acute light stimulation in the adult mPFC. (**a**) Representative pairwise correlations of L2/3 single units during acute ramp light stimulation (473 nm, 3 s) for a control (left) and ES mouse (right) at different developmental stages. (**b**) Average cumulative density functions of pairwise correlations during acute ramp light stimulation for control and ES mice at P11-12 (control n=11 recordings, 11 mice, ES n=10 recordings, 10 mice), P23-25 (control n=13 recordings, 6 mice, ES n=14 recordings, 5 mice) and P38-40 (control n=12 recordings, 5 mice, ES n=12 recordings, 5 mice). (**c**) Average intercept at 1^st^ and 3^rd^ quartile of correlation coefficients during acute ramp light stimulation for control and ES mice at different age. (Wilcoxon rank, P11-12, 1^st^ quartile p=0.385, 3^rd^ quartile p=0.162, LMEM, P23-25, 1^st^ quartile p=0.470, 3^rd^ quartile p=0.315, P38-40, 1^st^ quartile p=0.537, 3^rd^ quartile p=0.019). (**d**) Kolmogorov-Smirnov test score of the distance between pre and stim cumulative density function of correlation coefficients for control and ES mice. (Wilcoxon rank, P11-12, p=0.418, LMEM, P23-25, p=0.631, P38-40, p=0.033). Asterisks (* p<0.05, ** p<0.01, *** p<0.001) indicate significant differences (see Extended Data Tab. 1 for detailed statistics).

**Extended Data Fig. 7.**
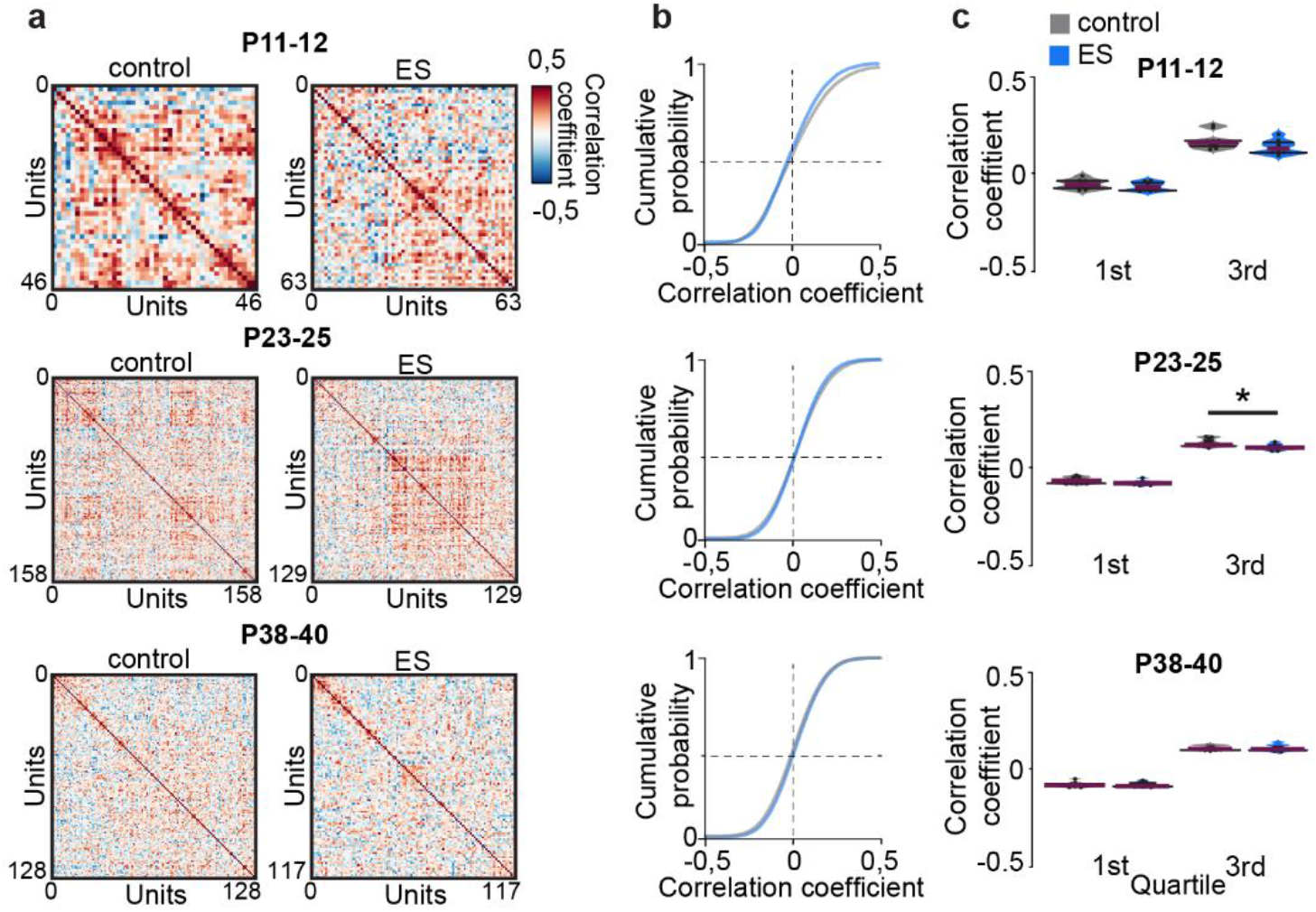
Transient early stimulation mildly affects network synchrony during spontaneous activity in the mPFC. (**a**) Representative pairwise correlations of L2/3 single units before acute ramp light stimulation for a control (left) and ES mouse (right) at different developmental stages. (**b**) Average cumulative density functions of pairwise correlations before acute ramp light stimulation for control and ES mice at P11-12 (control n=11 recordings, 11 mice, ES n=10 recordings, 10 mice), P23-25 (control n=13 recordings, 6 mice, ES n=14 recordings, 5 mice) and P38-40 (control n=12 recordings, 5 mice, ES n=12 recordings, 5 mice). (**c**) Average intercept at 1^st^ and 3^rd^ quartile of correlation coefficients before acute ramp light stimulation for control and ES mice at different age. (Wilcoxon rank, P11-12, 1^st^ quartile p=0.241, 3^rd^ quartile p=0.104, LMEM, P23-25, 1^st^ quartile p=0.100, 3^rd^ quartile p=0.036, P38-40, 1^st^ quartile p=0.970, 3^rd^ quartile p=0.911). Asterisks (* p<0.05, ** p<0.01, *** p<0.001) indicate significant differences (see Extended Data Tab. 1 for detailed statistics).

### Transient increase of neonatal activity alters excitation/inhibition balance in the adult mPFC

Network synchronization in gamma frequency results from interactions between excitatory and inhibitory units^36,37^. To elucidate the mechanisms of abnormal network synchronization upon stimulation in ES mice, we analyzed the response of individual units in L2/3 of the mPFC to acute ramp light stimulations. At P11-12, 25.3% of units in control mice and 20.3% of units in ES mice significantly increased their firing rate during ramp light stimulation. Only few units (control 0.9%, ES 3.4%) decreased their firing rates. In older mice, units with significantly increased (P23-25, control 31.2%, ES 32.7%; P38-40, control 33.7%, ES 25.5%) and decreased (P23-25, control 27.0%, ES 23.6%; P38-40, control 24.5%, ES 30.6%) firing rates were detected (Fig. 7a,b). The ratio of activated vs. inactivated neurons per mouse was similar across groups at P11-12 and P23-25, yet significantly reduced in ES mice at P38-40 compared to controls.

**Fig. 7.**
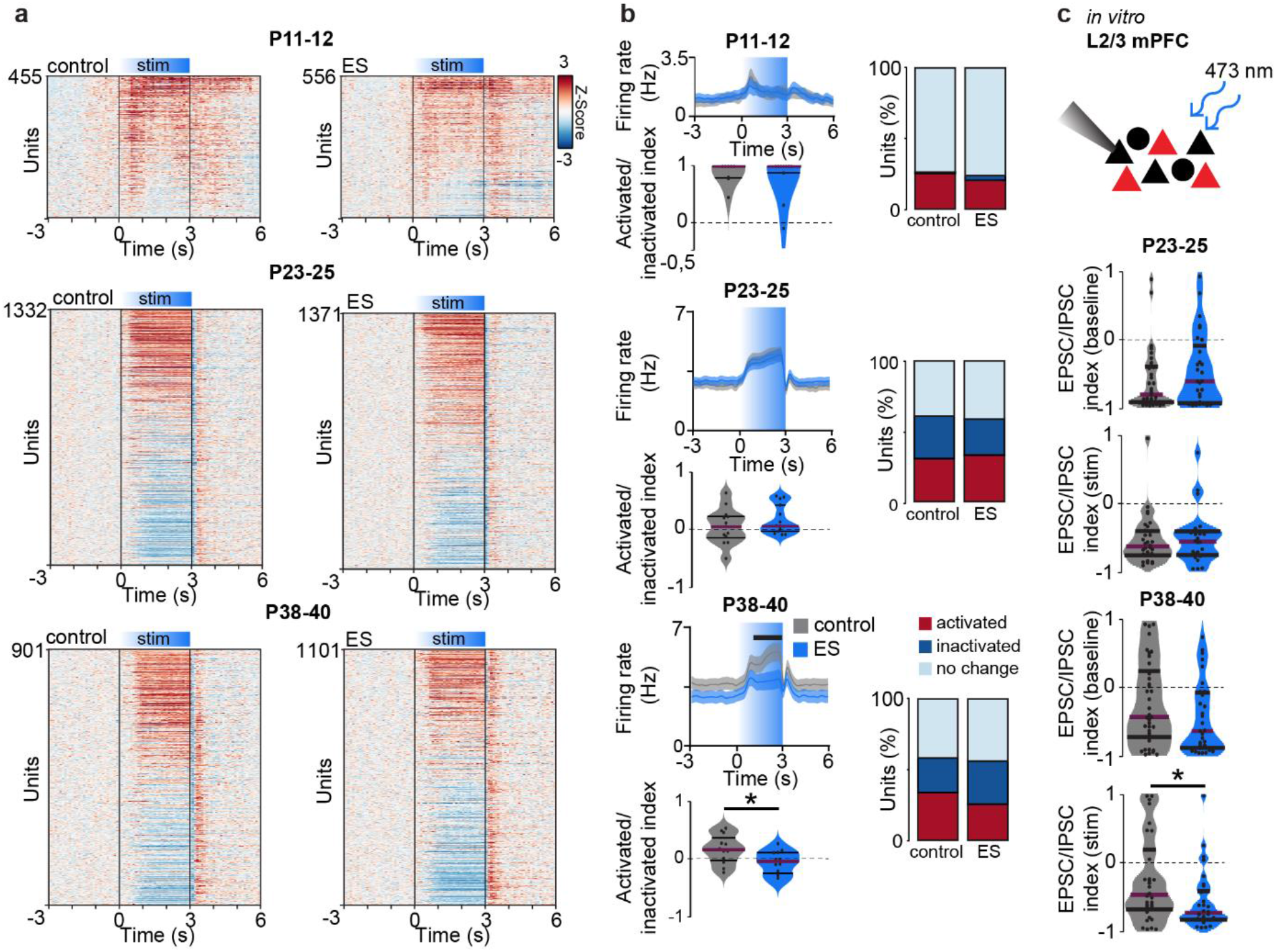
Transient early stimulation alters excitation/inhibition balance in the adult mPFC during acute light stimulation. (**a**) Single unit firing rates z-scored to pre-stimulation in response to acute ramp light stimulation (473 nm, 3 s) displayed for control (left) and ES mice (right) at P11-12 (control n=455 units, 11 mice, ES n=556 units, 10 mice), P23-25 (control n=1332 units, 6 mice, ES n=1371 units, 5 mice) and P38-40 (control n=901 units, 5 mice, ES n=1101 units, 5 mice). (**b**) Line plots displaying average firing rates during acute light stimulations (top left), violin plots showing the index of significantly activated vs. inactivated units (bottom left) and bar diagrams of the percentage of significantly activated and inactivated units for control and ES mice at P11-12, P23-25 and P38-40. (P11-12, LMEM, firing rate p<0.001, Wilcoxon rank, activated/inactivated index p=0.982, LMEM, P23-25, firing rate p=0.004, activated/inactivated index p=0.317, P38-40, firing rate p<0.001, activated/inactivated index p=0.033). (**c**) Top, schematic showing the protocol for in vitro whole-cell patch-clamp recordings from non-transfected L2/3 PYRs (black) during optogenetic stimulation of neighboring transfected cells (red) in the mPFC. Bottom, violin plots displaying EPSC/IPSC index during baseline and acute light stimulation (473 nm, square pulse, 1 s) for control and ES mice at P23-25 (control n=35 neurons, 5 mice, ES n=30 neurons, 5 mice) and P38-40 (control n=41 neurons, 6 mice, ES n=33 neurons, 4 mice). (LMEM, P23-25, baseline p=0.218, stim p=0.840, P38-40, baseline p=0.402, stim p=0.030). Black lines and asterisks (* p<0.05, ** p<0.01, *** p<0.001) indicate significant differences (see Extended Data Tab. 1 for detailed statistics).

Decreased gamma synchrony and stronger inhibition in P38-40 ES mice suggest that the transient increase of prefrontal activity at neonatal age causes long-term alterations of the balance between excitation and inhibition in the prefrontal circuitry. To test this hypothesis, we performed whole-cell patch-clamp recordings from non-transfected prefrontal L2/3 PYRs in coronal slices from control and ES mice. During acute light stimulation of ChR2(ET/TC)-transfected L2/3 PYRs (473 nm, square pulse, 1 s) the ratio of excitatory postsynaptic currents (EPSCs) to inhibitory postsynaptic currents (IPSCs) in non-transfected L2/3 PYRs was shifted towards inhibition for P38-40 ES mice compared to controls (Fig. 7c). In contrast, the ratio was similar between groups at younger age. Basic active and passive membrane properties as well as spontaneous inputs were not affected in ES mice (Fig. 7c, Extended Data Fig. 8).

**Extended Data Fig. 8.**
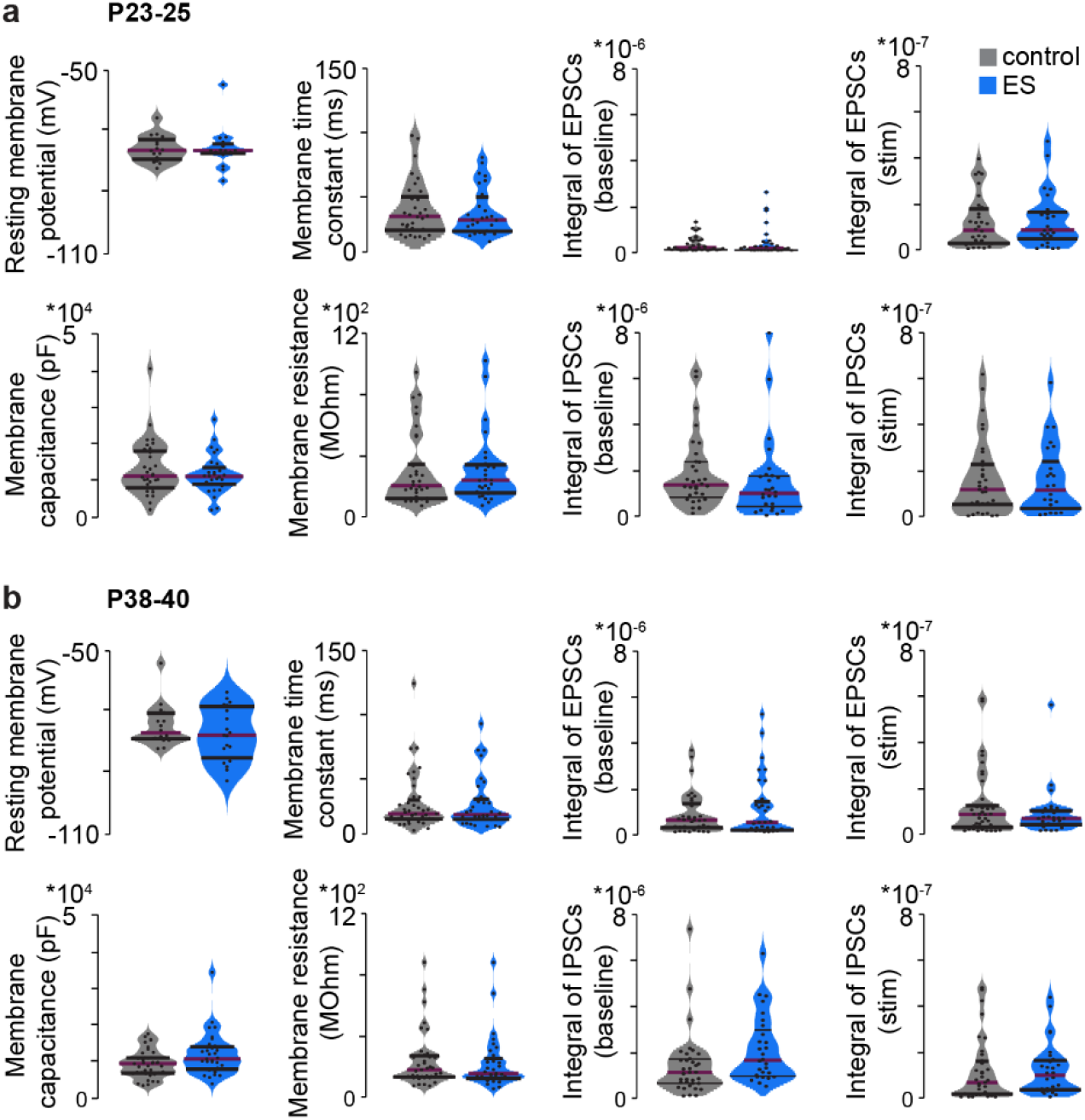
Transient early stimulation does not alter the passive and active membrane properties of non-transfected L2/3 PYRs. (**a**) Violin plots displaying passive and active membrane properties as well as properties of EPSCs and IPSCs induced by light stimulation in non-transfected L2/3 PYRs from control and ES mice at P23-25 (control n=35 neurons, 5 mice, ES n=30 neurons,5 mice). (LMEM, resting membrane potential p=0.545, membrane time constant p=0.426, EPSCs baseline p=0.743, EPSCs stim p=0.415 membrane capacitance p=0.218, membrane resistance p=0.564, IPSCs baseline p=0.234, IPSCs stim p=0.881). (**b**) Same as (a) for control and ES mice at P38-40 (control n=41 neurons, 6 mice, ES n=33 neurons, 4 mice). (LMEM, resting membrane potential p=0.526, membrane time constant p=0.907, EPSCs baseline p=0.339, EPSCs stim p=0.349 membrane capacitance p=0.304, membrane resistance p=0.436, IPSCs baseline p=0.332, IPSCs stim p=0.309). (See Extended Data Tab. 1 for detailed statistics).

Stronger inhibition might result from a higher survival rate of interneurons after transient activity increase during neonatal age^38^. To test this hypothesis, we performed immunohistochemical stainings for parvalbumin (PV) and somatostatin (SOM) and quantified the distribution of these two distinct subsets of inhibitory interneurons in the mPFC of control and ES mice during development. The density of SOM-positive neurons was significantly reduced, whereas the density of PV-positive neurons was significantly increased at P38-40 (Extended Data Fig. 9).

Fast-spiking (FS) PV-expressing interneurons that mature towards the end of the developmental period are critical for the generation of adult cortical gamma activity^36,39^. Therefore, the late emerging decrease of gamma synchrony in adult ES mice may result from disruption of these neurons. To test this hypothesis, we distinguished regular spiking (RS) and FS units in extracellular recordings from control and ES mice based on their spike waveform (Fig. 8a). This distinction revealed that the spontaneous firing rate of RS units is altered in ES mice at P23-25 and P38-40 compared to controls, whereas no changes were detected for FS units. In contrast, evoked activity during acute ramp light stimulation was reduced for RS and FS units in ES mice at P38-40, but normal earlier during development (Fig. 8c). Reduced evoked activity of FS units seems to be in opposition with the increased numbers of PV-positive neurons in adult ES mice. However, the FS firing rate is mainly reduced during the late phase of the ramp, whereas the initial peak is not altered. Taking into account that PV neurons inhibit pyramidal neurons but also other PV neurons^40^, we hypothesize that FS putatively PV neurons provide more potent inhibition and thereby reduce RS and FS firing rates after initial activation in ES mice.

**Fig. 8.**
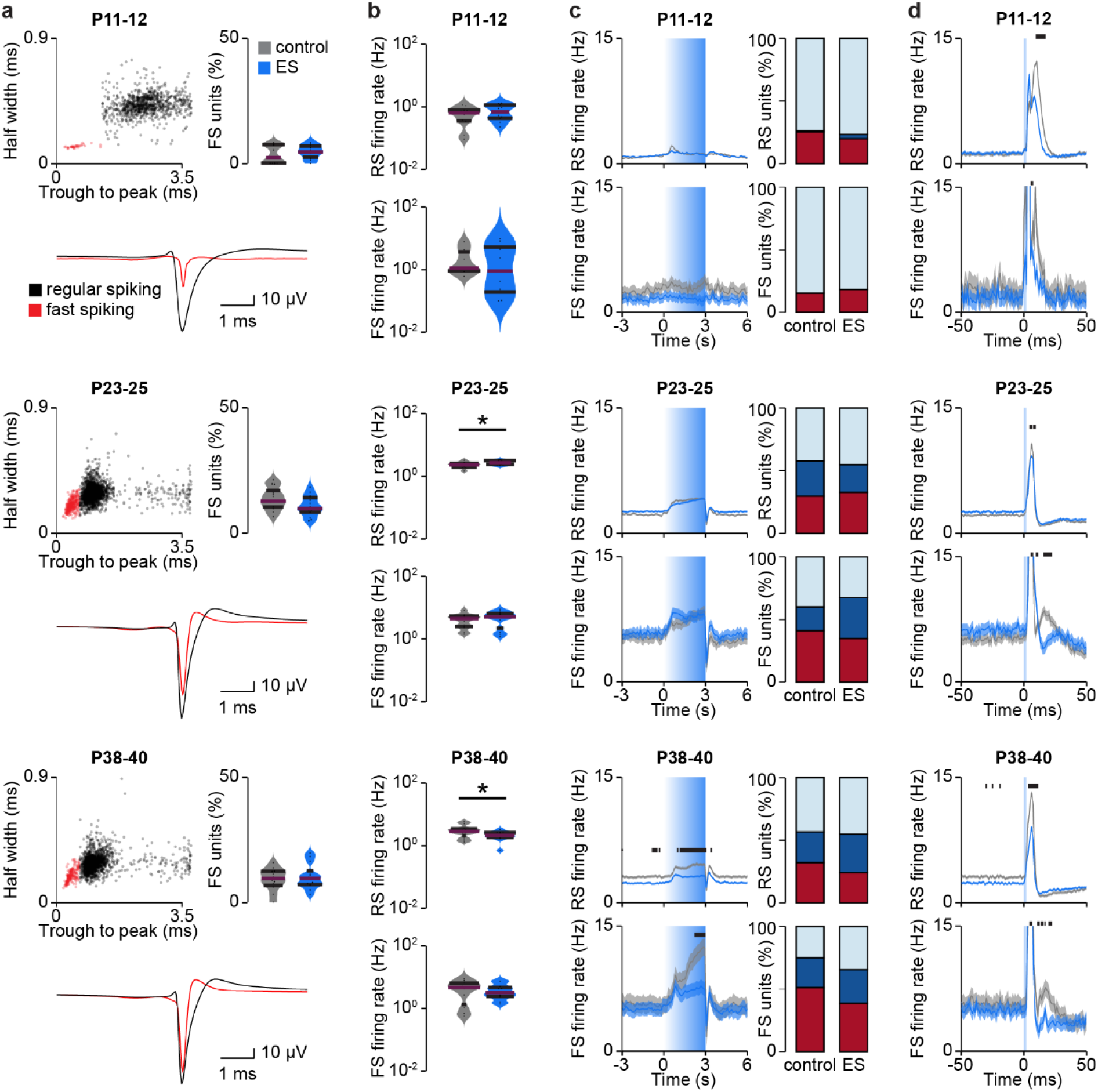
Transient early stimulation alters evoked inhibitory feedback from fast spiking units in the mPFC. (**a**) Scatter plots displaying half width and trough to peak duration (top left), average waveforms for RS and FS units (bottom), as well as percent of FS units for control and ES mice at P11-12 (control 428 RS and 13 FS units, 11 mice, ES 475 RS and 22 FS units, 10 mice), P23-25 (control 1140 RS and 185 FS units, 6 mice, ES 1220 RS and 141 FS units, 5 mice) and P38-40 (control 814 RS and 84 FS units, 5 mice, ES 992 RS and 104 FS units, 5 mice). (Wilcoxon rank, P11-12 p<0.500, LMEM, P23-25 p=0.114, P38-40 p=0.551). (**b**) Violin plots displaying spontaneous firing rate of RS and FS units for control and ES mice at P11-12, P23-25 and P38-40. (Wilcoxon rank, P11-12, RS firing rate p=0.418, FS firing rate p=0.680, LMEM, P23-25, RS firing rate p=0.020, FS firing rate p=0.357, P38-40, RS firing rate p=0.040, FS firing rate p=0.575). (**c**) Average firing rate during acute ramp light stimulation (473 nm, 3 s) and percent of significantly modulated units for control and ES mice at P11-12, P23-25 and P38-40. (LMEM, P11-12, RS firing rate p<0.001, FS firing rate p<0.001, P23-25, RS firing rate p<0.001, FS firing rate p<0.001, P38-40, RS firing rate p<0.001, FS firing rate p<0.001). (**d**) Average firing rate during acute pulse light stimulation (473 nm, 3 ms) for control and ES mice at P11-12, P23-25 and P38-40. (LMEM, P11-12, RS firing rate p<0.001, FS firing rate p<0.001, P23-25, RS firing rate p<0.001, FS firing rate p<0.001, P38-40, RS firing rate p<0.001, FS firing rate p<0.001). Black lines and asterisks (* p<0.05, ** p<0.01, *** p<0.001) indicate significant differences (see Extended Data Tab. 1 for detailed statistics).

Gamma synchronization in the adult cortex results from temporally coordinated excitatory drive and inhibitory feedback^37,39^. To investigate the timing of RS and FS firing, we performed acute stimulations with short light pulses of 3 ms duration. RS and FS units showed a pronounced peak in their firing rate for 5-10 ms in response to short light pulses (Fig. 8d). The similar peak time of RS and FS units indicates that the RS cluster contains a substantial number of non-transfected, indirectly activated units, in agreement with the sparse transfection achieved with IUE. FS units in control mice showed a second peak in their firing rate about 20 ms after the light pulse in P23-25 and P38-40 mice. The delay of 20 ms suggest the contribution of these units to gamma oscillations that have a typical cycle duration of 20 ms at 50 Hz. This second peak was significantly reduced in ES mice at P23-25 and P38-40. Of note, similar to ramp induced activity, the first peak was not affected for FS units, indicating that FS units provide stronger inhibition after initial activation in ES mice.

Thus, transiently increased activity at neonatal age alters the development of inhibitory feedback from FS interneurons and thereby, reduces evoked gamma synchronization of adult prefrontal circuits.

**Extended Data Fig. 9.**
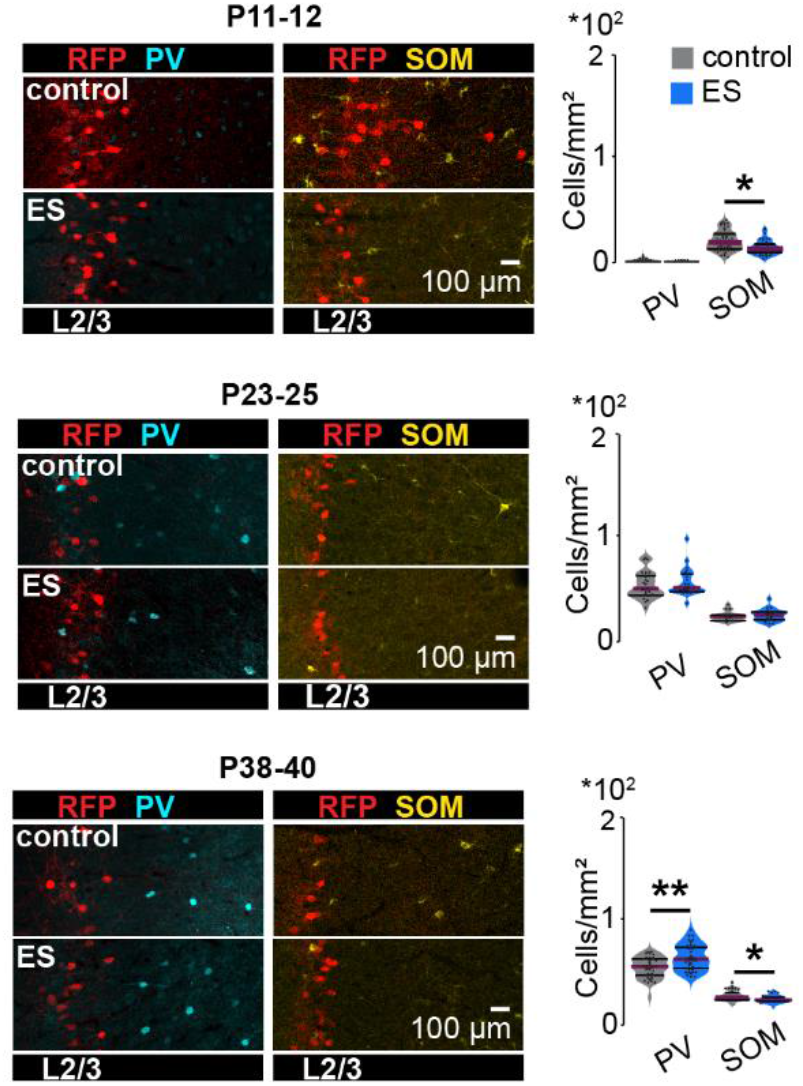
Transient early stimulation alters the density of SOM- and PV-expressing interneurons in the mPFC. Left, representative images showing PV and SOM immunostainings in the mPFC of control and ES mice at P11-12, P23-25 and P38-40. Right, violin plots displaying the density of PV-positive and SOM-positive neurons in L2/3 of the mPFC of control and ES mice at P11-12 (control: PV n=54 slices, 12 mice, SOM n=59 slices, 12 mice; ES, PV n=38 slices, 9 mice, SOM n=43 slices, 9 mice), P23-25 (control: PV n=25 slices, 5 mice, SOM n=25 slices, 5 mice; ES, PV n=27 slices, 6 mice, SOM n=25 slices, 6 mice) and P38-40 (control: PV n=36 slices, 9 mice, SOM n=36 slices, 9 mice; ES, PV n=40 slices, 10 mice, SOM n=43 slices,11 mice). (LMEM, P11-12, PV p=0.296, SOM p=0.044, P23-25, PV p=0.403, SOM p=0.390, P38-40, PV p=0.012, SOM p=0.012). Asterisks (* p<0.05, ** p<0.01, *** p<0.001) indicate significant differences (see Extended Data Tab. 1 for detailed statistics).

## Discussion

Seminal research identified electrical activity as a major contributor to the development of the mammalian cerebral cortex. Early activity influences neuronal migration, differentiation, and apoptosis^19–21^ as well as the establishment of synaptic connections^41,42^. Also in clinical settings, patterns of electroencephalographic activity of preterm infants provide prognostic value for neurodevelopmental outcome^43^. Several neuropsychiatric diseases have been proposed to be related to alterations in neuronal activity early in life^8,44^. However, fundamental questions still need to be addressed: how does electrical activity during early development impact adult cortical function? Does altered prefrontal activity during early development contribute to cognitive deficits later in life? Here, we address these questions and demonstrate that a transient increase of activity in the mouse mPFC during a short period of neonatal development critical for network formation causes long-lasting changes in inhibitory feedback and excitation/inhibition balance, leading to weaker evoked gamma band synchronization and ultimately, poorer cognitive abilities.

To manipulate developmental activity, we optogenetically stimulated the mPFC, inducing discontinuous activity patterns with similar dynamics as the ones spontaneously occurring. During early development the mammalian cortex shows discontinuous activity, with neuronal discharges organized in oscillatory rhythms alternating with electrically silent periods^45,46^. In the mPFC of neonatal mice, these 1-3 s-long oscillatory events with frequencies alternating between theta (4-12 Hz) and beta-low gamma (12-40 Hz) occur every 20-30 s^22,27^. The fast oscillations emerge as result of L2/3 PYRs activation^26^. Therefore, we used repeated ramp light stimulations (3 s duration, 6/min for 30 min) to activate L2/3 PYRs transfected with ChR2(ET/TC) by IUE and induced fast oscillatory discharges. At the age of transient early stimulation (P7-11), neurons have reached their final location in the cortical layers and are in the process of establishing synaptic connections^18,47^. Our stimulation protocol was designed to cause a modest increase of activity in the mPFC during this period critical for network formation. The stimulation not only augmented the level of activity but coordinated the prefrontal networks in fast oscillatory rhythms evolving from beta to gamma frequencies with age^35^. A causal link between this rhythmic organization and the long-term effects of transient early stimulation is still missing.

How does the transient increase of neuronal activity at neonatal age influence prefrontal development and ultimately behavior? The present data demonstrate that the early manipulation triggers a cascade of structural and functional changes in the mPFC leading to the impairment of cognitive abilities. On the morphological level, increased neonatal activity induced premature growth of dendrites in stimulated L2/3 PYRs. This is consistent with the activity-dependent growth of dendrites^48^ and reminiscent of the growth dynamics (i.e. initially excessive followed by arrested growth) during development in humans with autism spectrum disorders^49^. Activity of pyramidal neurons from P5 to P8 has been shown to regulate the survival of cortical interneurons^38,50^. Accordingly, we found an increased number of PV-expressing interneurons in ES mice. In contrast to previous studies^29,38,50^, this effect was specific for PV-expressing neurons, whereas the number of SOM-expressing neurons was reduced. Several differences in the experimental settings might explain this disparity: (1) Stimulation a few days later during developmental (P7-11 vs. P5-8) is expected to have a stronger effect on late maturing PV-expressing interneurons^29^; (2) Increased activity of a subset of pyramidal neurons (L2/3 PYRs vs. all PYRs) might cause different activation of interneuron subtypes; (3) Optogenetic (vs. chemogenetic) stimulation triggering fast oscillatory network activity might specifically engage PV-expressing interneurons.

Premature growth of dendrites likely affects the connectivity of stimulated neurons. Together with altered interneuron numbers, these structural changes led to a shift in the excitation/inhibition balance in the mPFC of ES mice towards inhibition. In addition to the general increase in inhibition, the timing of FS, presumably PV-expressing interneurons, was altered. In juvenile and adult control mice, brief activation of L2/3 PYRs induced a sharp peak in the firing rate of FS interneurons followed by a second peak about 20 ms later. This second peak, supposedly critical for synchronization in gamma frequency, was absent in ES mice. Accordingly, the transient increase of neuronal activity at neonatal age led to impaired synchronization of the prefrontal network in gamma frequency in young adults. This is consistent with the importance of PV-expressing FS interneurons for the generation of cortical gamma activity^37,39^. The late maturation of PV-expressing interneurons^29^ and gamma activity in the mPFC^35^ most likely underlie the delayed onset of these physiological effects. Of note, these effects were only evident during evoked activity, whereas spontaneous activity in the mPFC was largely normal, reflecting the moderate effects of stimulation protocol. This is consistent with alterations in evoked activity related to the early emergence of sensory symptoms in humans with autism spectrum disorders^44^.

Abnormal FS interneuron development impairs prefrontal gamma activity and cognitive flexibility in adults^51^. Accordingly, transient increase of neuronal activity at neonatal age ultimately resulted in impaired cognitive abilities in juvenile and young adult mice. Gamma activity in prefrontal L2/3 is particularly important for the maintenance of information during working memory tasks^52^. This is consistent with the specific impairment of ES mice in short-term memory and working memory tasks, as well as reduced social preference.

In conclusion, these data demonstrate that altered neuronal activity during early development induces structural and functional changes in the mPFC, ultimately resulting in impaired cognitive abilities. Even though cognitive symptoms are not the core deficits, they represent a devastating burden in neuropsychiatric diseases^3–5^. Altered cortical excitation/inhibition balance and impaired gamma activity are critical for cognitive dysfunctions^53–55^. Thus, altered developmental activity of cortical circuits might actively contribute to cognitive symptoms in neuropsychiatric diseases^14–18^.

Furthermore, the mechanisms described here might explain cognitive difficulties of preterm born humans experiencing excessive sensory stimulation in neonatal intensive care units (NICUs) (frequent handling associated with medical or nursing care, excessive noise and light levels) at a comparable stage of brain development (2nd-3rd gestational trimester)^56^. These stressful stimuli might trigger premature neuronal activity, perturbing the activity-dependent maturation of cortical networks^57^. Frontal regions are particularly vulnerable to conditions in NICUs ^58^ Correspondingly, preterm children are highly prone to frontally confined impairment, such as memory and attention deficits^59^. Thus, our findings lend experimental proof to the concept that neuronal activity during early development accounts for adult cortical function and cognitive performance, playing a critical role in neurodevelopmental and neuropsychiatric diseases^12,14,17^.

## Methods

### Animals

All experiments were performed in compliance with the German laws and the guidelines of the European Community for the use of animals in research and were approved by the local ethical committee (G132/12, G17/015, N18/015). Experiments were carried out on C57Bl/6J mice of both sexes. Timed-pregnant mice from the animal facility of the University Medical Center Hamburg-Eppendorf were housed individually at a 12 h light/12 h dark cycle and were given access to water and food ad libitum. The day of vaginal plug detection was considered E0.5, the day of birth was considered P0.

### In utero electroporation

Pregnant mice received additional wet food daily, supplemented with 2-4 drops Metacam (0.5 mg/ml, Boehringer-Ingelheim, Germany) one day before until two days after IUE. At E15.5, pregnant mice were injected subcutaneously with buprenorphine (0.05 mg/kg body weight) 30 min before surgery. Surgery was performed under isoflurane anesthesia (induction 5%, maintenance 3.5%) on a heating blanket. Eyes were covered with eye ointment and pain reflexes and breathing were monitored to assess anesthesia depth. Uterine horns were exposed and moistened with warm sterile phosphate-buffered saline (PBS). 0.75-1.25 μl of opsin- and fluorophore-encoding plasmid (pAAV-CAG-ChR2(E123T/T159C)-2A-tDimer2, 1.25 μg/μl) purified with NucleoBond (Macherey-Nagel, Germany) in sterile PBS with 0.1% fast green dye was injected in the right lateral ventricle of each embryo using pulled borosilicate glass capillaries. Electroporation tweezer paddles of 5 mm diameter were oriented at a rough 20° leftward angle from the midline of the head and a rough 10° downward angle from the anterior to posterior axis to transfect precursor cells of medial prefrontal L2/3 PYRs with 5 electroporation pulses (35 V, 50 ms, 950 ms interval, CU21EX, BEX, Japan). Uterine horns were placed back into the abdominal cavity that was filled with warm sterile PBS. Abdominal muscles and skin were sutured with absorbable and non-absorbable suture thread, respectively. After recovery from anesthesia, mice were returned to their home cage, placed half on a heating blanket for two days after surgery. Fluorophore expression in pups was detected at P2 with a portable fluorescence flashlight (Nightsea, MA, USA) through the intact skin and skull and confirmed in brain slices postmortem.

### Transient early stimulation

A stimulation window was made at P7 for chronic transcranial optogenetic stimulation in mice transfected by in utero electroporation. Mice were placed on a heating blanket and anesthetized with isoflurane (5% induction, 2% maintenance). Breathing and pain reflexes were monitored to assess anesthesia depth. The skin above the skull was cut along the midline at the level of the mPFC and gently spread with forceps. The exposed skull was covered with transparent tissue adhesive (Surgibond, SMI, Belgium). Mice were returned to the dam in the home cage after recovery from anesthesia. From P7-11 mice were stimulated daily under isoflurane anesthesia (5% induction, 2% maintenance) with ramp stimulations of linearly increasing light power (473 nm wavelength, 3 s duration, 7 s interval, 180 repetitions, 30 min total duration). Light stimulation was performed using an Arduino uno (Arduino, Italy) controlled laser system (Omicron, Austria) coupled to a 200 μm diameter light fiber (Thorlabs, NJ, USA) positioned directly above the tissue adhesive window. Light power attenuation was set to reach 10 mW in the brain, adjusted for measured light attenuation by the tissue adhesive (~30%) and the immature skull (~25%). Control animals were treated identical but stimulated with light of 594 nm wavelength that does not activate the expressed opsin ChR2(ET/TC).

### Electrophysiology and optogenetics in vivo

#### Acute extracellular recordings

Multi-site extracellular recordings were performed unilaterally or bilaterally in the mPFC of non-anesthetized and anesthetized P7-40 mice. Under isoflurane anesthesia (induction: 5%; maintenance: 2.5%), a craniotomy was performed above the mPFC (0.5 mm anterior to bregma, 0.1-0.5 mm lateral to the midline). Mice were head-fixed into a stereotaxic apparatus using two plastic bars mounted on the nasal and occipital bones with dental cement. Multi-site electrodes (NeuroNexus, MI, USA) were inserted into the mPFC (four-shank, A4×4 recording sites, 100 μm spacing, 125 μm shank distance, 1.8-2.0 mm deep). A silver wire was inserted into the cerebellum and served as ground and reference. Pups were allowed to recover for 30 min prior to recordings. For recordings in anesthetized mice, urethane (1 mg/g body weight) was injected intraperitoneally prior to the surgery. Extracellular signals were band-pass filtered (0.1-9,000 Hz) and digitized (32 kHz) with a multichannel extracellular amplifier (Digital Lynx SX; Neuralynx, Bozeman, MO, USA). Electrode position was confirmed in brain slices postmortem.

#### Chronic extracellular recordings

Multi-site extracellular recordings were performed in the mPFC of P23-25 and P38-40 mice. Under isoflurane anesthesia (5% induction, 2.5% maintenance), a metal head-post for head fixation (Luigs and Neumann, Germany) was implanted at least 5 days before recordings. Above the mPFC (0.5-2.0 mm anterior to bregma, 0.1-0.5 mm right to the midline) a craniotomy was performed and protected by a customized synthetic window. A silver wire was implanted between skull and brain tissue above the cerebellum and served as ground and reference. 0.5% bupivacaine / 1% lidocaine was locally applied to cutting edges. After recovery from anesthesia, mice were returned to their home cage. Mice were allowed to recover from the surgery, accustomed to head-fixation and trained to run on a custom-made spinning disc. For recordings, craniotomies were uncovered and a multi-site electrode (NeuroNexus, MI, USA) was inserted into the mPFC (one-shank, A1×16 recording sites, 100 μm spacing, 2.0 mm deep). Extracellular signals were band-pass filtered (0.1-9000 Hz) and digitized (32 kHz) with a multichannel extracellular amplifier (Digital Lynx SX; Neuralynx, Bozeman, MO, USA). Electrode position was confirmed in brain slices postmortem.

#### Acute light stimulation

Ramp (i.e. linearly increasing light power) light stimulation was performed using an Arduino uno (Arduino, Italy) controlled laser system (473 nm / 594 nm wavelength, Omicron, Austria) coupled to a 50 μm (4 shank electrodes) or 105 μm (1 shank electrodes) diameter light fiber (Thorlabs, NJ, USA) glued to the multisite electrodes, ending 200 μm above the top recording site. Acute stimulations were repeated 30 times.

### Electrophysiology and optogenetics in vitro

#### Patch-clamp recordings

Whole-cell patch-clamp recordings were performed from tDimer-negative L2/3 PYRs in the mPFC of P23–25 and P38-40 mice. Under anesthesia, mice were decapitated, brains were removed and sectioned coronally at 300 mm in ice-cold oxygenated high sucrose-based artificial cerebral spinal fluid (ACSF) (in mM: 228 sucrose, 2.5 KCl, 1 NaH2PO4, 26.2 NaHCO3, 11 glucose, 7 MgSO4; 310 mOsm). Slices were incubated in oxygenated ACSF (in mM: 119 NaCl, 2.5 KCl, 1 NaH2PO4, 26.2 NaHCO3, 11 glucose, 1.3 MgSO4; 310 mOsm) at 37°C for 45 min before cooling to room temperature. Slices were superfused with oxygenated ACSF in the recording chamber. Neurons were patched under optical control using pulled borosilicate glass capillaries (tip resistance of 3-7 MΩ) filled with pipette solution (in mM: 130 D-glucononic acid 49-53%, 130 Cesium-OH 50%, 10 HEPES, 0.5 EGTA, 4 Mg-ATP, 0.3 Na2-GTP, 8 NaCl, 5 QX-314-Cl; 285 mOsm, pH 7.3). Data was acquired using PatchMaster (HEKA Elektronik, MA, USA). Capacitance artifacts and series resistance were minimized using the built-in circuitry of the patch-clamp amplifier (EPC 10; HEKA Elektronik, MA, USA). Responses of neurons were digitized at 10 kHz in voltage-clamp mode.

#### Light stimulation

Square light stimuli of 472 nm wavelength and 1 s duration were delivered with the pE-2 LED system (CoolLED, Andover, UK).

### Histology

P5-40 mice were anesthetized with 10% ketamine (aniMedica, Germanry) / 2% xylazine (WDT, Germany) in 0.9% NaCl (10 μg/g body weight, intraperitoneal) and transcardially perfused with 4% paraformaldehyde (Histofix, Carl Roth, Germany). Brains were removed and postfixed in 4% paraformaldehyde for 24 h. Brains were sectioned coronally with a vibratom at 50 μm for immunohistochemistry or 100 μm for examination of dendritic complexity.

#### Immunohistochemistry

Free-floating slices were permeabilized and blocked with PBS containing 0.8% Triton X-100 (Sigma-Aldrich, MO, USA), 5% normal bovine serum (Jackson Immuno Research, PA, USA) and 0.05% sodium azide. Slices were incubated over night with primary antibody rabbit-anti-Ca2+/calmodulin-dependent protein kinase II (1:200, #PA5-38239, Thermo Fisher, MA, USA; 1:500, #ab52476, Abcam, UK), rabbit-anti-parvalbumin (1:500, #ab11427, Abcam, UK) or rabbit-anti-somatostatin (1:250, #sc13099, Santa Cruz, CA, USA), followed by 2 h incubation with secondary antibody goat-anti-rabbit Alexa Fluor 488 (1:500, #A11008, Invitrogen-Thermo Fisher, MA, USA). Sections were transferred to glass slides and covered with Fluoromount (Sigma-Aldrich, MO, USA).

#### Cell quantification

Images of immunostainings and IUE-induced tDimer2 expression in the right mPFC were acquired on a confocal microscope (DM IRBE, Leica, Germany) using a 10x objective (numerical aperture 0.3). tDimer2-positive and immunopositive cells were automatically quantified with custom-written algorithms in ImageJ environment. The region of interest (ROI) was manually defined over L2/3 of the mPFC. Image contrast was enhanced before applying a median filter. Local background was subtracted to reduce background noise and images were binarized and segmented using the watershed function. Counting was done after detecting the neurons with the extended maxima function of the MorphoLibJ plugin.

#### Dendritic complexity

Image stacks of tDimer2-positive neurons were acquired on a confocal microscope (LSN700, Zeiss, Germany) using a 40x objective. Stacks of 6 neurons per animal were acquired as 2048×2048 pixel images (voxel size 156*156*500 nm). Dendritic complexity was quantified by Sholl analysis in ImageJ environment. Images were binarized using auto threshold function and the dendrites were traced using the semi-automatic simple neurite tracer plugin. The geometric center was identified, and the traced dendritic tree was analyzed with the Sholl analysis plugin.

### Behavior

Mice were handled and adapted to the investigation room two days prior to behavioral examination. Arenas and objects were cleaned with 0.1% acetic acid before each trial. Animals were tracked online using video Mot2 software (Video Mot2, TSE Systems GmbH, Germany) or offline using the python-based tracking system ezTrack^60^.

#### Developmental milestones

Somatic and reflex development was examined every third day in P2-20 mice. Weight, body length, and tail length were measured. Grasping reflex was assessed by touching front paws with a toothpick. Vibrissa placing was measured as head movement in response to gently touching the vibrissa with a toothpick. Auditory startle was assessed in response to finger snapping. The days of pinnae detachment and eye opening were monitored. Surface righting was measured as time to turn around after being positioned on the back (max 30 s). Cliff avoidance was measured as time until withdrawing after being positioned with forepaws and snout over an elevated edge (max 30 s). Bar holding was measured as time hanging on a toothpick grasped with the forepaws (max 10 s).

#### Open field

At P16, Mice were positioned in the center of a circular arena (34 cm in diameter) and allowed to explore for 10 min. Behavior was quantified as discrimination index of time spent in the center and the border of the arena ((time in surround - time in center) / (time in surround + time in center)), grooming time, average velocity and number of rearing, wall rearing and jumping.

#### Object recognition

Novel object recognition (NOR, P17), object location recognition (OLR, P18) and recency recognition (RR, P21) were performed in the same arena as the open field examination. Mouse center, tail and snout position were tracked automatically. Object interaction was defined as the snout being within <1 cm distance from an object. For NOR, each mouse explored two identical objects for 10 min during the sample phase. After a delay period of 5 min in a break box, the mouse was placed back in the arena for the test phase, where one of the objects was replaced by a novel object. Behavior was quantified as discrimination index of time spent interacting with the novel and familiar object ((time novel object – time familiar object) / (time novel object + time familiar object)). OLR was performed similarly, but one object was relocated for the test phase instead of being exchanged. For RR, each mouse explored two identical objects during the first sample phase for 10 min, followed by a delay phase of 30 min, and a second sample phase of 10 min with two novel identical objects. After a second break of 5 min, the interaction time with an object of the first sample phase (old) and an object from the second sample phase (recent) was assessed during the test phase for 2 min. Behavior was quantified as discrimination index of time spent interacting with the novel and familiar object ((time old object - time recent object) / (time old object + time recent object)).

#### Maternal interaction

Maternal interaction was performed at P21 in the same arena as the open field examination. Two plastic containers were added to the arena, one empty and one containing the dam of the investigated mouse. Small holes in the containers allowed the mouse and the dam to interact. Behavior was quantified as discrimination index of time spent interacting with the empty container and the container containing the dam ((time dam container – time empty container) / (time dam container + time empty container)).

#### Spatial working memory

At P36-38, mice were positioned in the center of an elevated 8-arm radial maze. 4 arms contained a food pellet at the distal end (baited). On the first day, mice were allowed to examine the maze for 20 min or until all arms were visited. During the following 10 trials (2 trials on day 1 and 4 trials on day 2 and 3), mice were allowed to examine the maze until all baited arms were visited (for max 20 min) and arm entries were assessed. Visit of a non-baited arm was considered as reference memory error, repeated visit of the same arm in one trial as working memory error.

#### Spontaneous alteration

At P39, each mouse was positioned in the start arm of an elevated Y-maze. Visited arms during free exploration were monitored for 10 min. Percentage of alternations was calculated as (number of alternations / (entries - 2)). The test was used as habituation for delayed non-match-to-sample task.

#### Delayed non-match-to-sample task

At P39-40, mice were positioned in the start arm of an elevated Y-maze with access to the other arms containing a food pellet. After entering one arm, a central door was closed (sample choice). After the food pellet was consumed the mice were placed in the start arm for a second run (test choice) after a 30 s break. Each mouse performed 6 trials / day. Test choice was considered correct when visiting the arm not explored during sample phase.

### Data analysis

Data from in vivo and in vitro recordings were analyzed with custom-written algorithms in Matlab environment. In vivo data were band-pass filtered (500-9000 Hz for spike analysis or 1-100 Hz for LFP) using a third-order Butterworth filter forward and backward to preserve phase information before down-sampling to analyze LFP. For in vitro data, all potentials were corrected for liquid junction potentials (−10 mV). The resting membrane potential was measured immediately after obtaining the whole-cell configuration. To assess input resistance and membrane properties, 600 ms long hyperpolarizing current pulses were applied.

#### Power spectral density

For power spectral density analysis, 2 s-long windows of LFP signal were concatenated and the power was calculated using Welch’s method with non-overlapping windows. Spectra were multiplied with squared frequency.

#### Imaginary coherence

The imaginary part of complex coherence, which is insensitive to volume conduction, was calculated by taking the absolute value of the imaginary component of the normalized cross-spectrum.

#### Modulation index

For optogenetic stimulations, modulation index was calculated as (value stimulation - value pre stimulation) / (value stimulation + value pre stimulation).

#### Peak frequency and strength

Peak frequency and peak strength were calculated for the most prominent peak in the spectrum defined by the product of peak amplitude, peak half width and peak prominence.

#### Single unit analysis

Single unit activity (SUA) was detected and clustered using klusta^61^ and manually curated using phy (https://github.com/cortex-lab/phy). Modulation index of SUA firing rate was calculated on 3 s long windows pre- and during stimulation. Significance level was set at p<0.01 and calculated using Wilcoxon signed rank test for zero median for single stimulation trials. Single unit autocorrelation histogram was calculated using 0.5 ms bins followed by frequency spectrum computation using discrete Fourier transform. Oscillation score was calculated by dividing peak magnitude of detected peak frequency by average spectrum magnitude for pre- and during stimulation periods. For pairwise neuronal correlation SUA spike trains were convolved using a gaussian window with a standard deviation of 20 ms. Correlation of convolved spike trains was computed using Spearman’s rho. Cumulative distribution functions from before and during stimulations were compared using the two-sample Kolmogorov-Smirnov test. RS and FS units were distinguished by manually setting a threshold based on spike half width and trough-to-peak duration (FS, P11-12 halfwidth<0.31 ms, trough-to-peak<1.28 ms, P23-25 and P38-40 halfwidth<0.31 ms, trough-to-peak<0.64 ms).

#### EPSCs and IPSCs extraction

Voltage-clamp traces were demeaned and detrended with a median filter (mdefilt1). Traces were then deconvolved using a double exponential kernel using the OASIS toolbox (https://github.com/zhoupc/OASIS_matlab)^62^. After manual optimization of two separate kernels for EPSCs and IPSCs, the software was run with the “foopsi” model and a regularization parameter “lambda” set at the value of 10^-11^. The parameters “smin” and “b” were automatically optimized, separately for each trace. The deconvolved traces were then used to compute the integral of EPSCs and IPSCs for baseline and stimulation periods.

#### Statistics

Statistical analyses were performed in the Matlab environment or in R Statistical Software (Foundation for Statistical Computing, Austria). Data are presented as median ± median absolute deviation (MAD). Data were tested for significant differences (*p<0.05, **p<0.01 and ***p<0.001) using non-parametric Wilcoxon rank sum test for unpaired and Wilcoxon signed rank test for paired data or Kruskal-Wallis test with Bonferroni corrected post hoc analysis or Fisher’s exact test for binary data analysis. Nested data were analyzed with linear mixed-effect models with animal as fixed effect and Turkey multi comparison correction for post hoc analysis. See Extended Data Tab. 1 for detailed statistics.

## Notes

### Competing Interest Statement

The authors have declared no competing interest.

### Summary of Updates

While the previous version of the manuscript addressed the fundamental question whether electrical activity during early development impacts adult cortical function and ultimately behavior, the current version demonstrates how this happens. In the previous version, we concluded that altered developmental activity causes long-lasting impairments in prefrontal gamma synchrony and cognitive abilities. The new version of the manuscript shows the cascade of events caused by increased cortical activity during early development leading to cognitive impairments in young adults. Thereby this study links main concepts implicated with neuropsychological diseases (changes in early activity, altered inhibition by parvalbumin interneurons, changes in excitation/inhibition balance, impaired gamma synchrony and cognitive impairments) and brings them into a developmental perspective.

